# Dynamic flexibility of the murine gut microbiota to morphine disturbance enables escape from the stable dysbiosis associated with addiction-like behavior

**DOI:** 10.1101/2025.06.01.657215

**Authors:** Izabella Sall, Randi Foxall, Lindsey Felth, Anirudh Gaur, Jennifer L. Whistler, Cheryl A. Whistler

**Affiliations:** Department of Molecular, Cellular, & Biomedical Sciences, University of New Hampshire, Durham, NH, USA; Graduate program in Molecular and Evolutionary Systems Biology, University of New Hampshire, Durham, NH, USA; Center for Neuroscience, University of California–Davis, Davis, CA, USA; Department of Physiology and Membrane Biology, UC Davis School of Medicine, Davis, CA, USA

**Author notes:** Address correspondence to Cheryl A. Whistler, Jennifer L. Whistler.

**Keywords:** Microbiota, opioid, network analysis, compulsive behaviors, dysbiosis, opioid use disorder, morphine

## Abstract

Although opioids are effective analgesics, they can lead to problematic drug use behaviors that underlie opioid use disorder (OUD). Opioids also drive gut microbiota dysbiosis which is linked to altered opioid responses tied to OUD. To interrogate the role of the gut microbiota in a mouse model of OUD, we used a longitudinal paradigm of voluntary oral morphine self-administration to capture multiple facets of drug seeking and preserve both individual behavioral response and gut microbiota variation to examine associations between these two variables. After prolonged morphine consumption, only a subset of mice transitioned to a state we define statistically as compulsive. In compulsive mice, morphine fragmented the microbiota networks which subsequently reorganized to form robust novel connections. In contrast, the communities of non-compulsive mice also changed but were highly interconnected during morphine disturbance and maintained more continuity post morphine suggesting dynamic flexibility. Compulsive mice displayed a greater loss of functional diversity and a shift towards a new stable state dominated by potential pathobionts, whereas non-compulsive mice better preserved genera associated with gut health and broader functional diversity. These findings highlight how persistent and stable gut microbiota dysbiosis aligns with long-term behavioral changes underlying OUD, potentially contributing to relapse.

## INTRODUCTION

Opioids are the gold standard for acute and chronic pain management, but their prolonged use can lead to treatment limiting side-effects that cascade into the development of an opioid use disorder (OUD) defined by the *Diagnostic and Statistical Manual of Mental Disorders* (DSM-5^TM^) as a problematic pattern of opioid use leading to clinically significant impairment or distress^1-4^. OUD is diagnosed by a constellation of physiological and behavioral abnormalities with individuals presenting at least two of eleven criteria^4^. Diagnosis of an OUD does not require manifestation of all criteria, and not all affected individuals necessarily display the same core criteria. This underscores the complex phenotype of OUD and the possibility that variability stems from distinct but parallel biological processes that have yet to be fully elucidated. Rodent models have provided valuable insights into the biological effects of prolonged morphine exposure that may contribute to OUD, although they only partially capture the outlined DSM-5 diagnostic criteria^5-14^. For example, criteria such as awareness of a problem or the desire to cut down^4, 5^ cannot be modeled in mice, whereas tolerance, withdrawal, craving, and increased motivation to obtain opioids can be modeled, with the caveat that behavioral data can be limited by its subjective nature, though this can be improved by using multiple tests and measures of related behaviors. Even though mice can display variability in modelled behaviors, including morphine-induced tolerance and compulsive drug-seeking^15, 16^, paralleling the heterogeneity displayed by humans, most studies are designed to intentionally limit within-treatment variability to strengthen statistical inferences. Consequently, using extreme experimental treatments that drive homogeneity of responses erases informative variation that could be used to better decipher correlations, and eliminate confounders.

In addition to their intended analgesic effects, opioids cause unwanted side effects by binding with opioid receptors in peripheral neurons of the gastrointestinal tract, which disrupt neural mechanisms essential for maintaining normal gastrointestinal function and homeostasis^17-19^. This in turn, causes microbiota dysbiosis^6, 7, 9, 15, 20-24^, reflecting an imbalance in microbial community composition or loss of functions, likely through changes in the gastrointestinal environment, such as increased intestinal permeability or a “leaky gut”, reduced chloride secretion resulting in increased fluid and electrolyte absorption, and decreased transit (i.e. constipation)^17-19^. Morphine-induced microbiota dysbiosis has, consequently, been linked to a variety of opioid-induced biological and behavioral changes—some aligning with OUD diagnostic criteria--including analgesic tolerance^6-8^, dependence^9-12^, and opioid-seeking^13, 14^, whereas other changes linked to the microbiota, even if not used as criteria, may share etiology with OUD behaviors or increase the risk of OUD (e.g. opioid reward^21, 23, 25-27^ and hyperalgesia^21, 28^). However, as a common feature of morphine exposure, microbiota dysbiosis alone cannot explain why mice do not universally display these drug-induced behavioral changes when exposed to clinically relevant drug doses. By preserving natural variability in response to morphine and controlling for common morphine-induced microbiota changes, we recently showed functional preservation of butyrate production, which is redundant among diverse taxa, was naturally protective for the development of opioid tolerance^15^. This highlights how not all morphine-driven changes that correlate with OUD outcomes necessarily drive OUD diagnostic traits and the value of using natural variation to help decipher general or side effects from informative associations, or where underlying mechanisms of behaviors may differ.

Opioids induce dramatic shifts in microbiota composition^6, 7, 9, 15, 20-24^, raising questions about the capacity of the gut microbiota to recover from disruption, known as resilience^29, 30^ and where it cannot, and what impact this has on OUD progression. Whereas defining a “healthy” gut microbiota remains controversial, it is widely held that diversity (the membership and distribution of taxa and/or functions) and complexity (the ecological interactions among taxa) are key indicators of microbiota homeostasis^31^, contributing to resiliency and stability of the microbiota in response to perturbations^29, 30, 32^. In contrast, a reduction of diversity, stability, and resilience from baseline are hallmarks of dysbiosis which can contribute to disease progression and recurrence^29, 30^. It is notable that while some behavioral changes that align with OUD diagnostic criteria resolve rapidly upon drug removal (e.g. antinociceptive tolerance and physical dependence)^6, 7, 15^, others persist even after cessation of drug (e.g., relapse to drug seeking)^16, 33^. The persisting effects of opioid use that can drive relapse are arguably the greatest obstacle for recovery in those with OUD and the least understood. Furthermore, the extent to which microbiota dysbiosis persists beyond drug cessation, and whether it contributes to long lasting behavioral change, has not been experimentally addressed. When disturbed, the microbiota can transition to alternative stable states—distinct community structures potentially with altered functionality or signaling^30, 34-36^. Disturbances can either be continuous (“press”) that lead to a stable yet altered assemblage, or short-term (“pulse”), in some instances allowing the microbiota to recover to its original assemblage^30^. Considering whether changes of the gut microbiota during and after prolonged morphine disturbance contributes to drug seeking behavior and relapse could help direct future mechanistic studies and elucidate targets for OUD prevention.

Here, we expanded upon our prior study of morphine-induced antinociceptive tolerance and the microbiota changes during 18-weeks of opioid self-administration^15^ to more comprehensively evaluate whether the microbiota could play a role in complex, variable and persisting behavioral changes reflecting transition to compulsive-like opioid-seeking. We used an extended three-phase paradigm that models aspects of OUD in mice, including escalation of drug-seeking, failure to extinguish drug-seeking as a model of craving, and reinstatement after prolonged abstinence as a measure of relapse, in an effort to align with multiple DSM-5 criteria^15,16^ to identify microbiota signatures associated with OUD progression. In compulsive mice, morphine dramatically disrupted the microbiota community network structure which convergently transitioned to a new stable and morphine adapted state, with a greater increase in pathobionts, and lower functional diversity relative to non-compulsive mice reflective of persistent dysbiosis. In contrast, the microbiota of non-compulsive mice, while also disrupted and reaching an alternative stable state, displayed more complex connectivity during morphine disturbance and when morphine was removed the microbiota regained more normal variability, and networks displayed greater continuity with the pre-morphine state. Relatively higher abundances of recognized beneficial commensals, and greater functional diversity were also preserved in the microbiota of non-compulsive mice indicating the alternative stable state of their microbiota, though less resistant to future morphine disturbances of diversity, was more functionally resilient through its inherent variability.

## MATERIALS AND METHODS

### Animal Use and Care

Age-matched male (n = 20; 16 purchased from Jackson Laboratory and 4 bred in-house at UC Davis) and female (n = 4; 4 bred in-house at UC Davis) wildtype C57/BL6J mice, *Mus musculus*, were subjected to the oral morphine self-administration paradigm in two separate cohorts (n = 22, 8-weeks-old upon entering lever training with 0.2% saccharin reward and 11-weeks-old at first morphine exposure), as previously described^16^, and were fed Tekland 2018 diet (Inotiv, USA) which was formulated with soybean meal containing 150-340 mg/kg isoflavone. Briefly, we combined aspects of both voluntary home cage morphine consumption and operant lever pressing for drug in a model that provides for significant morphine exposure (home cage consumption), and the ability to model motivation for drug (lever pressing). Mice underwent operant training to light and tone cues to and learned to press a lever to receive a saccharin reward using Med Associates operant conditioning chambers (Fairfax, VT). Mice who successfully completed operant training with saccharin (Sigma-Aldrich, St. Louis, MO) were entered into this study. Mice were individually housed with access to two bottles, one with water and one with morphine sulfate (MS) (Mallinckrodt Pharmaceuticals, St. Louis, MO) + 0.2% saccharin to improve palatability and their operant responding to drug 5 days a week. After two days with access only to water, mice were placed in the operant chambers and their lever pressing behavior recorded. The paradigm is three phases: 1) Operant Self-Administration of drug, 2) Extinction, and 3) Reinstatement. During the self-administration phase, morphine self-administration is tracked both in the home cage and in the weekly operant sessions for 18 weeks. The concentration of MS in the home cage gradually escalated from 0.3 mg/mL week 1, to 0.5 mg/mL week 2, and then to a final concentration of 0.75 mg/mL for the remainder of the paradigm. In the Extinction phase, morphine is removed from the home cage and mice are placed in the operant chamber every day for up to 12 days with cues but no morphine reward. Mice were considered extinguished when they pressed less than 20% of their normal responding as measured in their final operant session at week 18 or after 12 consecutive days. After extinction, mice experienced abstinence when they were returned to their home cage for two weeks with access only to water and no morphine, and where no behaviors were monitored or recorded. Finally, in the Reinstatement phase, mice were returned to the operant chamber, given the cue and a single non-contingent morphine reward after which no lever presses produced morphine to assess whether they return to pressing the lever.

All mice had food *ad libitum* and were provided running wheels for extra enrichment. Individual housing was necessary to monitor morphine consumption and changes in drug-seeking behaviors for each mouse. All procedures involving animals were reviewed and approved by the Institutional Animal Care and Use Committee of the University of California, Davis (IACUC protocol number: 22085).

### Calculation of Compulsivity Scores

Measures of drug seeking during each phase of the behavioral paradigm were compiled into a sub-score, and these were then combined into an overall composite compulsivity score to categorize individual mice as either “compulsive” or “non-compulsive”, as previously described^16^. Briefly, behavioral measures taken during weeks 16-18 of the self-administration phase as well as each of the other phases of the paradigm were each separately Z-scored and summed to create phase-specific sub-scores for operant drug seeking, extinction of drug seeking, and reinstatement of drug seeking. For the final composite compulsivity score, the three behavioral sub-scores (self-administration, extinction, and reinstatement) were combined into a final composite score. Mice were then categorized based on their interquartile standard deviation (IQD) from the population mean: those 1 IQD above the mean were classified as compulsive, a term we use merely as a vernacular tool to describe their categorization into statistically defined groups, and those that were below this 1 IQD threshold as non-compulsive. Mice scoring 2 IQD above or below the population mean were further classified as the “most” compulsive (i.e., highest composite compulsivity scores) and “least” compulsive (i.e., lowest composite compulsivity scores) within the cohorts.

### Statistical Analysis of Animal Data

Linear regression analyses using Pearson’s correlation coefficient determined whether compulsivity significantly correlated with total morphine consumption, preference for morphine over saccharin, or morphine antinociception using the R package ggpubr (v 0.6.0)^37^. Normality of data was assessed using a Shapiro-Wilk test and homogeneity of variance was assessed using Levene’s test. Data with a p-value greater than 0.05 was considered normal. Welch Two-Sample T-test was used to compare differences between groups where assumptions for normality and homogeneity of variance were met. The Wilcoxon Rank Sum test was used when assumptions for normality and homogeneity of variance were not met. All statistics were conducted using the R package stats (v 4.3.0)^38^. Visuals were generated using R 4.2.0 software^38^, BioRender, and some composite figures were modified with Adobe Illustrator.

### 16S rDNA Library Preparation and Sequencing

DNA extraction, 16S rDNA library preparation, and sequencing of feces collected throughout the paradigm were performed as previously published, and the raw 16S amplicon sequencing reads were made publicly available in NCBI under the accession number PRJNA1098090^15^. Briefly, microbial genomic DNA was extracted from frozen feces; concentrations were standardized for two-step PCR amplification. The V4-V5 16S regions were amplified using 515F and 926R primers^39, 40^ modified to include the forward and reverse primer pad and linker sequences, and amplicons indexed using xGen UDI Primer Pairs. Indexed amplicons were pooled and purified prior to sequencing on the Illumina NovaSeq 6000 platform (2 x 250 bp paired end) at the Hubbard Center for Genome Studies at the University of New Hampshire (Durham, NH). Computations were performed on Premise, a central, shared HPC cluster at the University of New Hampshire (Durham, NH) supported by the Research Computing Center and PIs who have contributed compute nodes.

### Processing of 16S rDNA Sequencing Reads

16S rDNA sequencing reads were processed as previously described^15^. Briefly, demultiplexed raw sequencing reads were processed using the cutadapt plugin^41^ in the QIIME 2 2020.2 pipeline^42^, R package DADA2 (v1.18)^43, 44^, and custom scripts for artifact removal to generate a quality-controlled dataset. A taxonomic filter was applied to remove unclassified phyla, chloroplasts, and mitochondria, along with an additional prevalence filter to remove dataset-wide singletons with the expectation that they were erroneously produced (i.e., sequencing error), thereby maintaining true singletons within individual samples. Taxonomy was assigned using the GreenGenes reference database (v13.8)^45^ and a maximum likelihood phylogenetic tree with bootstraps was estimated from MAFFT (v7.305b)^46^ alignments using RAxML (v8.2.10)^47^. All data components (i.e., taxonomic assignments, experiment metadata, counts, sequences, and a phylogenetic tree) were stored together using the R package phyloseq (v1.40.0)^48^ for downstream analyses. Subsequent analyses utilized amplicon sequence variant (ASV) datasets that were either agglomerated by phylogeny (similar ASVs into one representative taxon) or taxonomy (genus level) using phyloseq (v1.40.0)^48^. All analyses and visuals were produced using R Studio (v4.2.0)^38^. For a summary of 16S processing and analysis, see Supplemental Figure 1.

Overall gut microbiota β-diversity (community composition differences) and temporal changes assessed by Bray-Curtis dissimilarity (relative abundance) was analyzed using principle coordinates analysis (PCoA) in phyloseq (v1.40.0)^48^. Permutation Multivariate Analysis of Variance (PERMANOVA)^49^ tests were performed to statistically evaluate the gut microbiota βdiversity (Bray-Curtis dissimilarity, all ASVs) of individual mice each week over the course of the morphine self-administration phase using the pairwise.adonis2 function in the R package pairwiseAdonis^50^. Significant differences in microbiota compositional diversity relative to week one of morphine self-administration were detected and persisted through week 18. For analyses of associations with early and late self-administration microbiota, two overlapping datasets were generated into an early stage (weeks one to 12) and late stage (weeks eight to 18). A four-week overlap between these two stages (weeks eight to 12) accounts for independent cohort treatment and uneven collection of feces across the two cohorts, as this design allowed for temporally more even representation of samples (Supplemental Table 1).

### Analysis of Microbiota Stability, Variability, and Resilience

To visualize microbiota compositional shifts in response to morphine disturbance and determine whether they were alternative and stable following the conclusion of the morphine self-administration phase, β-diversity at the ASV-level was initially assessed using Jensen-Shannon dissimilarity, which quantifies the similarity between taxa abundance distributions^51^, and analyzed using PCoA^34^. PERMANOVA^49^ tests determined any statistical differences between pre- and post-paradigm and compulsivity using the pairwise.adonis2 function in the R package pairwiseAdonis (v 0.4.1)^50^. The effect of cohort was also accounted for using the following formula: Jensen-Shannon dissimilarity ∼ Compulsivity Classification with strata = Cohort to control for variability introduced by cohorts (i.e., controlling for cohort as a batch effect) (Supplemental Table 2). Prior to using PERMANOVA, pairwise permutation tests^52^ for homogeneity of multivariate dispersion^53^ determined any statistical difference in group variances of compulsivity throughout the paradigm using the betadisper and permutest functions in the R package vegan (v2.6-4)^54^ (Supplemental Table 2). To determine the number of distinct microbiota states, Gaussian mixture modeling (GMM) was applied to the PCoA axis one coordinates using the R package the mclust (v6.1.1)^55^. GMM fits the PCoA axis one coordinates as a mixture of multiple Gaussian (normal) distributions, with each distribution representing a cluster, which corresponds to a potential microbiota state^55^. The modelName = “V” argument was used to allow variance to be estimated separately for each identified cluster^55^. The optimal number of clusters was selected using Bayesian Information Criterion (BIC), and model selection was validated using bootstrap sequential Likelihood Ratio Testing (LRT)^55^.

To assess stability of the alternative state, overall β-dispersion at the ASV level using Jensen-Shannon dissimilarity was computed using the betadisper function in vegan (v2.6-4)^54^ pre- and post-paradigm. Pairwise permutation tests^52^ for homogeneity of multivariate dispersion^53^ determined any statistical differences between pre- and post-paradigm using the permutest function in the R package vegan (v2.6-4)^54^. Additionally, we tested whether the standard deviations (SD) of individual ASVs differed significantly between samples taken during pre- and post-paradigm. We employed statistical tests to compare variance distributions using R packages dplyr (v1.1.4)^56^ and ggplot2 (v3.5.1)^37^. A dataset consisting of ASV-specific SD values for the two conditions was prepared from and OTU abundance table using the R package dplyr to group the ASVs by experience and calculate the SD values for each ASV. The SD for each ASV at pre- and post-paradigm was plotted using ggplot2. The F-test was applied to compare the SD distributions of the two conditions under the assumption of normality.

Temporal changes of α-diversity were characterized using observed diversity (ASV richness) and Simpson’s Diversity Index during each phase of the paradigm. For analysis of variability in within- and between-mouse α-diversity within a phase or changes in variability across phases, normality was first assessed using the Shapiro-Wilk test in the R package stats (v 4.3.0)^38^. When normality assumptions were met, Barlett’s test was used to determine significant differences in between-animal variability (population variance of α-diversity) and the F-test was used to determine significant differences in within-animal variability (SE distributions of α-diversity) using the R package stats (v 4.3.0)^38^.

A relative quantitative measure of resilience^57, 58^ was used to determine whether the gut microbiota of compulsive and non-compulsive mice changed differently or showed different trajectories of “recovery” (resiliency) following morphine press and pulse disturbances. The resilience index (RL) was calculated for observed richness based on ASV counts using the estimate_richness function in phyloseq (v 1.40.0)^48^. The RL index formula was^57, 58^:

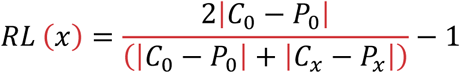

In this formula, C_0_ and P_0_ represent ASV richness using a reference state and a time when the microbiota is disturbed, respectively, while C_x_ and P_x_ represent ASV richness of a reference state and a post-disturbance (recovery) state, respectively. ASV richness of the reference state was averaged across all mice in the population, while ASV richness of the disturbed (P_0_) and recovered (P_x_) states was averaged across samples within each individual mouse. For this study, C_0_ was equal to C_x_. RL is bounded by -1 and +1, with +1 indicating full recovery, a, 0 indicating no recovery from the disturbed state post-disturbance, and –1 indicating no change during disturbance, but a change post-disturbance. To assess the normal patterns of variability that do not reflect systematic disturbance, an RL was calculated using three samples pre-paradigm where no systemic disturbance occurred, providing a threshold to distinguish changes in diversity greater than normal variability, thereby limiting false positives.

During continuous morphine exposure during the self-administration phase (press disturbance), reference, disturbed, and recovery states of observed diversity (ASV richness) corresponded to pre-paradigm, morphine self-administration, and extinction, respectively, where the reference corresponded to the population average, and the disturbed and recovered states were average diversity per each individual mouse. An RL index was also calculated from pulse disturbance that occurred during reinstatement where the reference, disturbed, and recovery states corresponded to extinction, reinstatement, and post-paradigm, respectively, where the reference diversity is the population average, and where disturbed and recovered represent one microbiota sample per individual. Differences in RL scores for ASV richness between compulsive and non-compulsive mice were assessed using the Welch Two-Sample T-test if assumptions of normality and homogeneity of variance were met, or the Wilcoxon Rank Sum test when assumptions of normality and homogeneity of variance were not met in the R stats^38^.

### Microbiota Association Networks

Microbial association networks inferred from the gut microbiomes of mice before entering the 18-week morphine self-administration phase (pre-paradigm), during the morphine self-administration phase, and after the morphine self-administration phase concluded (post-paradigm) were compared between non-compulsive and compulsive gut microbiomes. Community members were grouped at the genus level using the tax_glom function from phyloseq (v1.40.0)^48^. The top seventy out of 184 taxa grouped at the genus level were at the highest frequency and used to measure co-occurrence networks. SPRING associations were measured in the gut microbiota split by whether they were taken from not compulsive or compulsive mice. Microbial association networks for each group were inferred using the signed distance, nlambda, and rep.num (100) functions in the R package NetCoMi^59^. For reproducibility, microbial association network analyses used the random seed “12345”. Differential networks were inferred from SPRING association networks using fisher test adjusting for false discovery rate, although none were found.

### Analysis of Predicted Functional Profiles

Functional profiles of gut microbiota from compulsive and non-compulsive mice were predicted from representative ASVs of 16S rDNA sequences using phylogenetic investigation of communities by reconstruction of unobserved states (PICRUSt 2.0) pipeline^60^. Gene content, represented by Enzyme Classification (EC) numbers, (i.e., gene family copy number of ASVs and abundances per sample) per ASV was predicted^61^ by aligning ASVs to reference sequences and placement into a reference phylogenetic tree^62-64^. MetaCyc functional pathways and associated abundances were inferred from EC number abundances^65^. Differential abundance testing of predicted functional pathways between tolerant and non-tolerant mice across the different paradigm phases was analyzed using the LinDA model in the R package ggpicrust2 (v1.7.3)^66^.

### Differential Abundance, Biomarker, and Indicator Species Analyses

Community members, grouped at the genus level, that were different in abundance between the “most” compulsive (determined by scores above the population average + 2 IQD) and the “least” compulsive mice (determined by scores below the population average - 2 IQD) across the phases of the paradigm were determined using the differential test within the R package corncob (v0.3.1)^67^. Non-normalized counts were used with the Wald (abundance) or LRT (variability) setting within the differential test to distinguish community members associated with the covariate of interest controlling only for sequencing run, cohort, and paradigm phase when power was sufficient. When a phase did not have sufficient power (i.e., sampling of feces less than three times per mouse at a given phase, including extinction, reinstatement, and post-paradigm), only the covariate of interest was specified to avoid overfitting the model. The count abundance for each genus was fit to a beta-binomial model using the logit link functions for both the mean and overdispersion simultaneously. The null and non-null overdispersion models were specified with the same confounding variables (paradigm experience when testing the entire dataset, and sequencing plate and cohort when testing within each paradigm phase) to identify only genera having differential abundances associated with the covariate of interest and were not confounded with other covariates. The list of differentially abundant and/or variable community members produced was further analyzed using linear discrimination analyses of effect size (lefser v1.14.0)^68, 69^ to identify any community members that most likely explain differences between a covariate of interest. Community members were visualized with the amp_heatmap function in the R package ampvis2 (v 2.8.9)^70^, using a log2 color scale and the minimum abundance threshold of 0.07. Additionally, we used indicator species analyses to determine community members unique, or characteristic to the most or least compulsive mice^71^. To do this, the abundance data was transformed into presence/absence (0 for not present, and 1 for present), and each community member assessed for indicator value of either compulsive or non-compulsive mice, and significance tested using 999 permutations in the R package indicspecies (v1.7.15)^71^.

### Correlations of Gut Microbiota with Compulsivity Score

Genera significantly associated with the degree of compulsivity (measured as the composite compulsivity score) were identified with the R package MaAsLin2 (v1.18.0)^72^. Individual genera were fit to linear regression models of center log-ratio normalized counts, with a minimum prevalence threshold set at 0.05 across all samples. For the global analysis—examining correlations with degree of compulsivity across all phases—fixed effects included composite compulsivity score, cohort, and fecal sample number (serving as a proxy for timepoint), whereas random effects included sequencing plate and individual mouse. For analyses within specific paradigm phases, fixed effects included composite compulsivity score, fecal sample number, and cohort, and random effects were limited to individual mouse when power was sufficient. When a phase did not have sufficient power (i.e., sampling of feces less than three times per mouse at a given phase, including extinction, reinstatement, and post-paradigm), only composite compulsivity scores and sample number were identified as fixed effects, and no random effects were specified as to avoid overfitting the model. The remaining MaAsLin2 execution parameters were set to the following: standardize = FALSE (since composite compulsivity scores were pre-standardized) and transform = NONE (as center log-ratio serves as both a normalization and transformation method). Volcano plots were generated to visualize significant features (plotted as -log10(q-value) versus correlation coefficient) with a significance threshold of q-value = 0.25, accounting for multiple comparisons and controlling for false positives. Only significant features are highlighted, with labels indicating the genus and corresponding ASV.

### Predictive Modelling

Supervised machine learning was conducted following the best practices^73^ using the R package mikropml (v1.6.1)^74^. L2 logistic regression and random forest models were trained on genus abundance counts to classify mice as compulsive or non-compulsive (categorical outcome) or to predict genera most strongly associated with compulsivity score (continuous outcome). Two predictive modeling algorithms (logistic regression versus random forest)^75^ were first assessed for whether a more interpretable (logistic regression, Supplemental Figure 2) or a higher performing (random forest, Supplemental Figure 3) algorithm was better suited to the dataset. Where logistic regression models the probability of an outcome by estimating the strength and direction of associations between predictive features (e.g., taxa abundances) and the outcome^76^, random forest builds multiple decision trees to identify the most important features (e.g., taxa abundances) contributing to model accuracy and performance^76^. Multiple metrics were used to assess model performance (Supplemental Figures 2 & 3, evaluation metrics described further below). Random forest models consistently outperformed L2 logistic regression models and thus were selected for analyses.

Random forest models were trained with data from each phase of the paradigm. Data was pre-processed by centering and scaling abundance counts, collapsing perfectly correlated genera, and removing genera with zero variance, before the data was split into training and testing sets. For weighted classification modeling (categorization), case weights were calculated based on the proportion of compulsive and non-compulsive mice to address the imbalance between these groups. Approximately 70% of the data was allocated to the training set when a model was generated for each phase of the paradigm. For regression modeling (composite compulsivity score), for each of 100 random seeds, data were randomly split into 80% training and 20% testing sets specifically for modeling using continuous data (i.e., composite compulsivity scores).

Model performance was evaluated using multiple metrics. For weighted classification modeling, the metrics included cross-validated AUC (training set) and test set AUC, balanced accuracy, F1 score, precision, and recall, as these are suitable for an imbalanced dataset^77, 78^. For regression modeling, cross-validated RMSE (training set), test set RMSE, and R-squared values were used for assessing model performance. A baseline RMSE model was calculated from the composite compulsivity score mean as a reference point for further evaluation of a model’s performance. Finally, feature importance was assessed using permutation testing. Following the collapse of perfectly correlated genera, each genus in the test data was randomly shuffled 100 times and a new permutation performance (AUROC) was calculated. Genera with a permutation AUROC significantly lower than the original test AUROC (p-value < 0.05) were considered important contributors to model performance. Genera with the greatest decrease in AUROC upon permutation were deemed most important to model performance.

## RESULTS

### Not all mice developed compulsive drug-seeking behaviors

To capture behavioral changes indicating the transition to compulsive-like drug-seeking, we applied a longitudinal paradigm with three phases—self-administration, extinction, and reinstatement—wherein behaviors that align with the DSM-5 diagnostic criteria for OUD related to the time and effort spent obtaining and using opioids were measured^4, 5, 16^ in two cohorts of wild-type mice (n=16 and 6) (Figure 1A-B). During the self-administration phase, mice had *ad libitum* access to both morphine and water in their home cages^16^ and weekly operant lever-pressing sessions to retrieve morphine, cued by light and sound, measured motivation for reward (Figure 1B, red gradient box). Increased lever pressing indicates increased motivation for drug, as mice are without home-cage morphine for two days prior to their lever pressing session. After assessment of tolerance to morphine antinociception^15^, mice entered the extinction phase and had daily timed sessions identical to their weekly drug sessions, but lever presses were not rewarded (morphine was never available) (Figure 1B, teal box). The number of days of unrewarded lever pressing before lever pressing declined to 20% of that on week 18 (determined on an individual basis) indicated the rate of drug-seeking. Following extinction, mice were returned to their home cage for a 2-week abstinence period during which only water was available in the home cage, and there were no operant sessions. After this abstinence period mice entered the final phase where they were assessed for reinstatement of drug-seeking. Mice were placed in the operant chamber and drug was both cued and delivered without the need to press a lever (a single non-contingent reward) after which the session mirrored the regular self-administration paradigm except no lever presses were rewarded (Figure 1B, yellow box). Importantly, all mice retrieved the reward, but shorter latencies for reward retrieval along with subsequent head port entries and lever presses to attempt access to additional rewards signifies reinstated motivation to obtain morphine, modelling “relapse”. At the study conclusion, recorded measures taken during each phase were compiled into sub-scores for each phase, where the self-administration phase was also separated into early and late stages for some analyses (e.g. Figure 1, Supplemental Figure 4). Final sub-scores included a single self-administration sub-score based on behaviors during the last 3 weeks of the self-administration phase, an extinction sub-score, and a reinstatement sub-score. In addition, these individual phase sub-scores were combined to generate an overall “compulsivity score” as described previously^16^ (Supplemental Figure 4). As a vernacular tool, mice were categorically classified as “compulsive” drug seekers if their score was >1 interquartile deviation (IQD) above the population mean and “non-compulsive” drug-seekers if their score was lower than this >1 IQD threshold (Figure 1C). Using this statistic, eight mice (36.3%) were categorized as compulsive (red circles) and 14 mice as non-compulsive (grey circles) (Figure 1C). Mice statistically categorized as compulsive diverged from non-compulsive mice in the degree of drug seeking late in the self-administration phase and remained significantly different during the reinstatement phase (Figure 1D). Though population distribution of individual mice by final composite compulsivity score was bimodal, mice with the most extreme composite scores do not fit smoothly within the populations to which they are assigned and some mice with intermediate scores due to moderate behaviors are categorized as non-compulsive despite a lack of absolute separation between the populations (Figure 1E).

**Figure 1:**
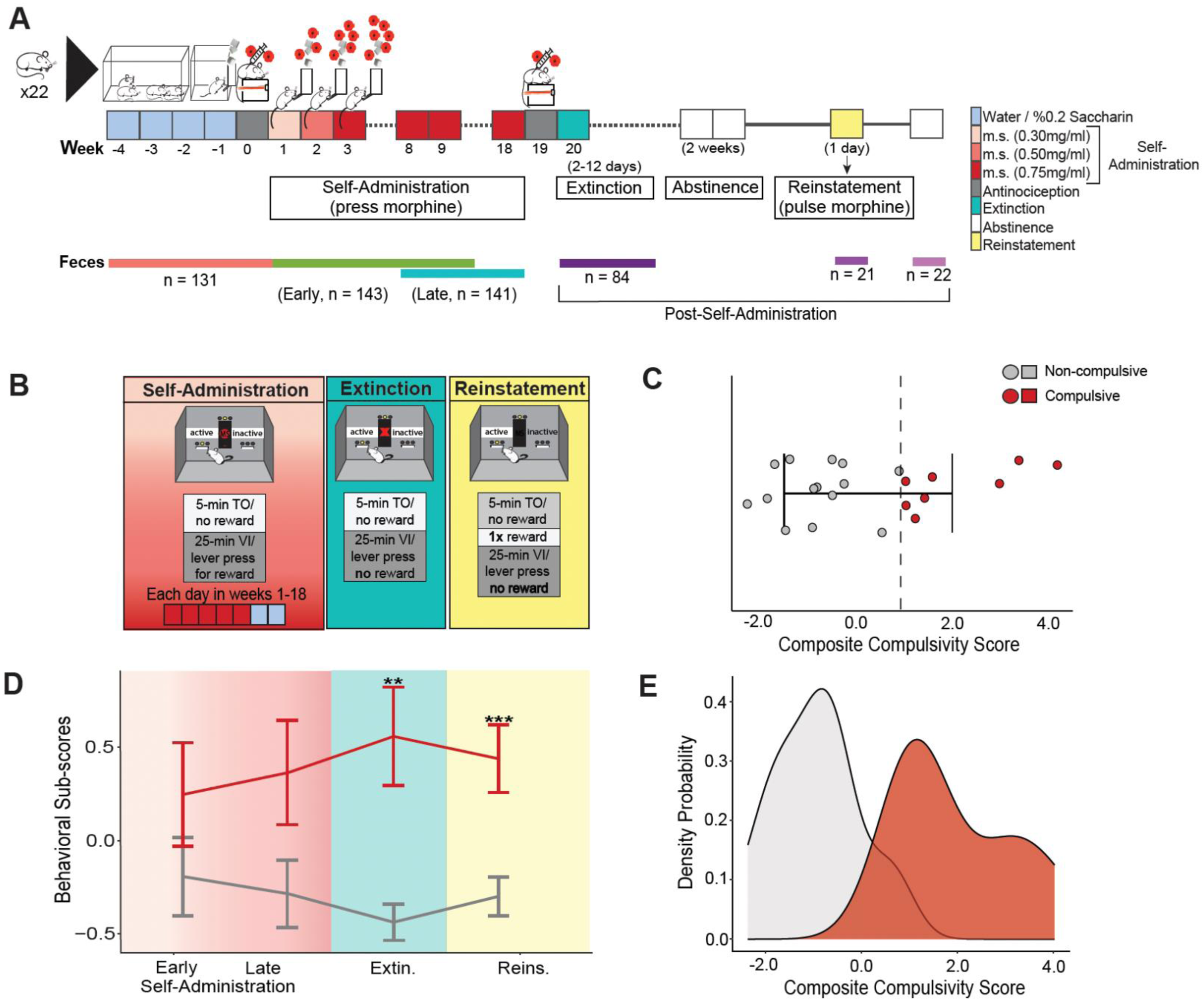
Wild-type mice diverge over time in their display of drug-seeking behaviors allowing assignment to compulsive or non-compulsive populations. A) Three phase animal behavioral paradigm for development of compulsive behaviors including a self-administration phase with increasing doses of morphine (0.3 mg/mL week one, 0.5 mg/mL week two, and 0.75 mg/mL weeks three to 18), an extinction phase where reward was withheld followed by a two-week abstinence period where only water was available in the home cage, and a reinstatement phase where reward was not contingent on lever-pressing. Voluntary morphine consumption, preference for morphine over saccharin, development of antinociceptive tolerance, and differences in drug-seeking behaviors were monitored. Feces were collected from mice (n = 22) throughout the study, color coded by phase, and used to generate 16S (V4-V5 region, see methods) sequencing reads for microbiota analyses. B) Weekly operant sessions (red gradient box) were used to document motivation to obtain drug by lever pressing to obtain reward. After the 18-week morphine self-administration phase, the rate and extent of extinction (teal box) and reinstatement (yellow box) were measured (see methods for more details). C) Mice were statistically categorized as compulsive if their composite compulsivity score is >1 interquartile deviation (IQD) above the population mean as indicated by the grey dashed line (red) or non-compulsive if below this threshold (grey). D) The trajectory of divergence of the two populations of mice were visualized using individual mouse behavioral sub-scores, which included an early self-administration sub-score using measures only from the first three weeks and a late self-administration score based on behaviors from weeks 16-18, the latter of which was used as the final sub-score for self-administration (see^16^). Statistical differences in sub-scores of compulsive (red line) and non-compulsive (grey line) mice were assessed using the Student’s t-test. Significance thresholds are indicated as ** = p < 0.01 and *** p < 0.001. E) Population distribution of individual mice categorized as compulsive (red) or non-compulsive (grey) visualized using a density probability plot, which estimates the probability distribution of composite scores as a continuous curve to identify modal patterns and overall spread in the data from observed data points.

As previously shown, individual compulsivity scores did not correlate with other behaviors related to morphine use and thought to predispose risk of OUD including quantity of morphine taken or preference for morphine over another reward (saccharin), reinforcing that these do not necessarily predict altered drug-seeking behaviors (Supplemental Figure 4A &, B). Furthermore, even though antinociceptive tolerance is an OUD-diagnostic criteria that can lead to an escalation of drug intake, degree of tolerance also did not correlate with compulsivity score (Supplemental Figure 4C), suggesting that biological mechanisms that underlie tolerance are not necessary for the development of more complex compulsive-like behaviors. This suggests that if the microbiota plays a role in the development of compulsive behaviors, different functions of the gut microbiota could contribute to compulsive-like behaviors than to antinociceptive tolerance.

### The alternative stable state of the microbiota of non-compulsive mice retained a relatively higher degree of variability than the microbiota of compulsive mice during and post-morphine

Compulsive-like behaviors used for statistical categorization only significantly diverged post-morphine self-administration (Figure 1D), suggesting persisting effects of morphine on microbiota composition could be related to differences in behaviors. We therefore visualized how the microbiota from individual mice categorized as either compulsive or non-compulsive (Figure 2A, dark closed circles and light closed circles respectively) was changed by morphine comparing pre-paradigm microbiota (weeks –2 to 0; orange) to microbiota collected after the self-administration phase, during extinction, reinstatement, and post-paradigm combined (week 19-onward, pink) via principle coordinates analyses (PCoA). This revealed that prolonged morphine exposure drove a microbiota compositional shift to an alternative community structure (Figure 2A, pink) that significantly differed from the diverse pre-paradigm microbiota community structure in both compulsive and non-compulsive mice alike (Figure 2A) (pPERMANOVA = 0.005; Figure 2A, Supplemental Table 2). Gaussian mixed models of the distribution of extracted PCoA axis one coordinates further indicate two significantly distinct populations (p-value = 0.001), reflective of the microbiota either prior to or after prolonged morphine. This aligned with previous findings that morphine strongly drives convergent changes of microbiota assemblages^15^ but further demonstrated that the alternative state persisted after morphine was removed. Whereas no other PCoA axis separated the microbiota of mice to discernible clusters corresponding to compulsive and non-compulsive groups, this is expected since nuanced but meaningful microbiome associations are largely masked by common morphine driven β-diversity shifts^15^.

**Figure 2:**
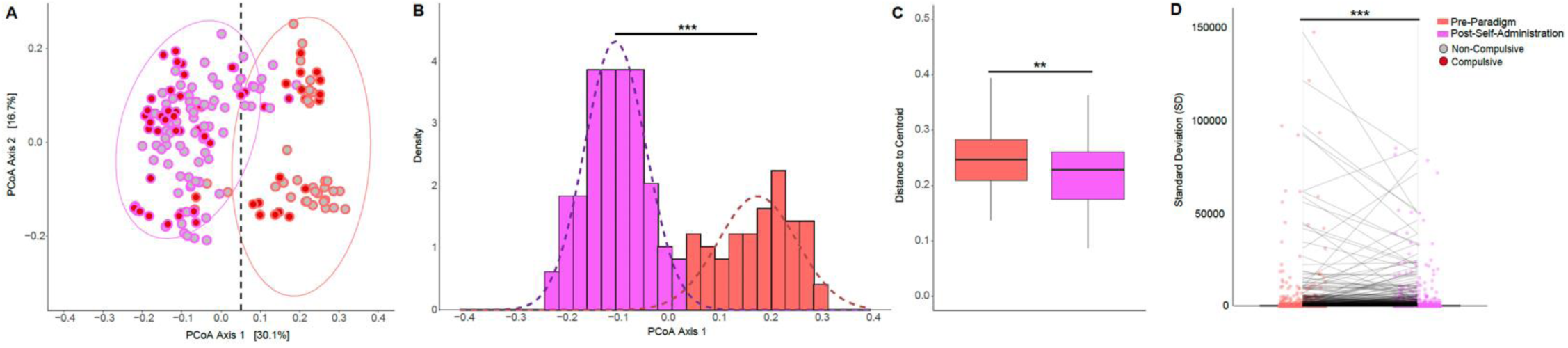
Gut microbiota of all mice shifted to an alternative stable state in response to prolonged morphine exposure. A) Temporal changes in the gut microbiota β-diversity (all ASVs, Jensen-Shannon dissimilarity) prior to (pre-paradigm, orange), and after (post-self-administration, pink), the 18-week morphine self-administration phase as visualized by principal coordinates analysis (PCoA). See Supplemental Table 2 for details. The black dashed line corresponds to the separation between the two normal distributions demonstrated in panel B. B) Determination of the number of clusters using Bayesian Information Criterion (BIC) in Gaussian mixture modeling (GMM) via the R package mclust (v6.1.1) as visualized by frequency distributions of PCoA axis one coordinates extracted from (A) as a measure of microbiota state(s). The number of clusters was statistically validated by bootstrap sequential Likelihood Ratio Testing (LRT)^55^ (*** = p < 0.001). C) Mean β-dispersion (microbiota variability) (All ASVs, Jensen-Shannon dissimilarity) of post-self-administration (pink) relative to pre-paradigm (orange). Differences in microbiota variability were evaluated using pairwise permutation tests, with significant levels denoted as follows: ** = p < 0.01. D) Comparison of standard deviations (SD) of all individual ASVs between samples taken during pre-paradigm (orange) and post-self-administration (pink). Differences in the SD distributions of all individual ASVs during pre-paradigm (orange) and post-self-administration (pink) were evaluated using the F-test under the assumption of normality, with significant levels denoted as follows: *** = p < 0.001.

Next, using variability as a proxy for stability, the spread (β-dispersion) of microbiota composition of all mice grouped by pre-paradigm or post-self-administration were compared (Figure 2C, D). On the whole population level, mean gut microbiota variability significantly decreased post-self-administration (p-value = 0.007) (Figure 2C). Individual ASV variability measured by standard deviations of their abundance across multiple samples also significantly decreased (p < 0.0005) post-self-administration compared to pre-paradigm (Figure 2D). This pattern of reduced microbiota variability at both global and individual ASV levels following prolonged morphine reinforces that mice with limited genetic variation, and from the same vivarium, can normally exhibit substantial variation in their microbiota and that the convergent, alternative state of the microbiota following prolonged morphine disturbance is stable and persisting (Figure 2C, D).

Another consideration for whether an alternative state is stable is its resistance to change and resiliency, or the ability of a microbiota to recover from disturbances^29, 30^. Though many factors can influence resiliency of a microbiota, it is widely held that diversity, particularly captured by α-diversity metrics such as richness (the number of unique species) and evenness (considering both number and relative abundance of different species), promotes resiliency. Subtle differences in the timing of morphine-driven changes in α-diversity, which can reflect progressing dysbiosis^29, 30, 79^, are linked to morphine-driven antinociceptive tolerance^15^. Therefore, we visualized the patterns of within-mouse microbiota disturbance and recovery from the “press” prolonged morphine exposure, as well as the subsequent “pulse“ exposure during reinstatement and evaluated whether patterns of changing diversity differed between compulsive and non-compulsive mice. Comparison of the average α-diversity in samples collected during press morphine disturbance with average diversity of pre-paradigm samples revealed ASV richness decreased (observed diversity or unique ASVs present) only in the microbiota of compulsive mice (on average a loss of 20 ASVs, or 3%). In contrast, ASV richness of non-compulsive mice generally varied by the same magnitude (20 ASVs), but without a net loss. Press morphine disturbance convergently decreased community evenness (Simpson’s Diversity Index) in both compulsive and non-compulsive mice and the magnitude of change was not different (on average an increase of ∼0.02 reflecting reduced diversity, p-value = 0.29). Combined, these data suggest morphine changes α-diversity and, at least at the population level, the resulting disturbance was not entirely convergent between compulsive and non-compulsive mice in terms of ASV richness.

We next visualized how changes in ASV richness within individual compulsive and non-compulsive mice differed from each other by comparing diversity patterns as mice transition through different phases of the paradigm (Figure 3A). For context, microbiota richness within individual mice is normally variable but not different between compulsive and non-compulsive mice pre-paradigm (F-test p-value = 0.94, Figure 3B), ∼290 ASVs from day to day (Figure 3A & B, pre-saccharin). This variability decreased modestly to ∼200 ASVs during operant training with saccharin (the reference condition for microbiota responses to morphine as all mice were trained on saccharin) which also caused a convergent shift in composition in all mice^15^ (Figure 3A). With press morphine disturbance, average ASV richness of all individual mice converged showing significantly less between-animal variability and also significantly less temporal within-animal variability (saccharin reference to self-administration, Figure 3A & B), ranging from only ∼50 ASVs in compulsive mice to ∼100 ASVs for non-compulsive mice where non-compulsive mice maintained significantly more within mouse variation (p-value = 0.0053, Figure 3B). Interestingly, through the extinction, reinstatement, and post-paradigm phases, the ASV richness of compulsive mice remained in this convergent state with no significant difference in variability between extinction and reinstatement, and no significant change in within-mouse temporal variability between self-administration and extinction (Figure 3A &B), suggesting the microbiota of compulsive mice, shaped by press morphine, resisted change as the paradigm progressed. In contrast, even though diversity of non-compulsive mice also did not significantly change or diverge during extinction, within-mouse variability did significantly increase during extinction (Figure 3A & B). Notably, the microbiota richness of individual non-compulsive mice gradually became more variable or divergent from one another, especially during reinstatement—when mice once again get a single pulse morphine dose (Figure 3A & B). This indicates that the microbiota composition established during press morphine in non-compulsive mice maintained a greater degree of within-mouse variability.

**Figure 3:**
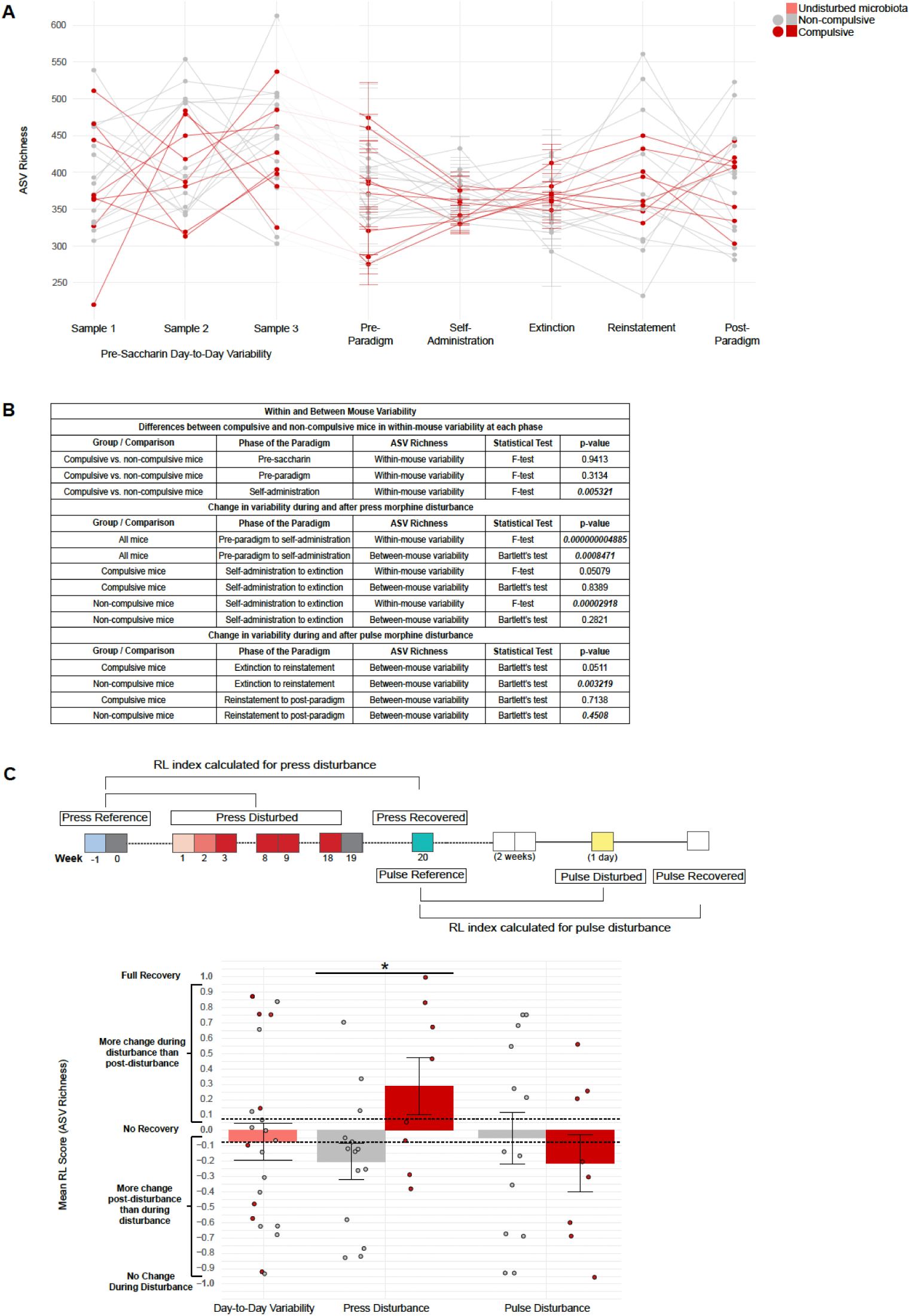
The morphine-adapted microbiota of compulsive mice was relatively more stable than the microbiota of non-compulsive mice. A) Temporal change of average ASV richness of individual compulsive (red) and non-compulsive (grey) mice through the three-phase behavioral paradigm. The first three timepoints (Sample 1, Sample 2, Sample 3) represent normal day-to-day variability of the gut microbiota in the absence of disturbance as determined from samples collected pre-saccharin (during weeks -4 to -3, Figure 1A). Samples collected during saccharin and the paradigm are aggregated and averaged by mouse where multiple samples were available (i.e., pre-paradigm, self-administration, extinction). See Figure 1A for paradigm details. B) When normality assumptions were met, differences in between-animal ASV richness variability were assessed using Bartlett’s test, whereas differences in within-animal ASV richness variability were assessed using the F-test. Comparisons were made for compulsive and non-compulsive mice within a phase, or across phases within compulsive mice or non-compulsive mice. C) Simplified visual of microbiota samples used to calculate the RL index during the press and pulse morphine disturbances based on the paradigm schematic in Figure 1A. The RL index was calculated as the change in ASV richness of the disturbed and recovered microbiota, each relative to the ASV richness of the reference microbiota. Normal day-to-day variability was visualized by calculating an RL index score using ASV richness (pink bar) to establish a threshold for normal variation (black dashed lines). RL scores for the recovery of ASV richness^57, 58^ in compulsive and non-compulsive mice following press or pulse morphine disturbances was calculated by averaging ASV richness across all mice for the reference state, while ASV richness was averaged across samples within individual mice for the disturbance and recovery states. Differences in mean RL scores between compulsive (red bar) and non-compulsive (grey bar) mice were evaluated using the Welch Two-Sample T-test or Wilcoxon Rank Sum test, with significant levels denoted as follows: * = p < 0.05.

Microbiota resiliency was next quantified using a resilience (RL) index for observed diversity (ASV richness) to further define responses and recovery of the microbiota in compulsive and non-compulsive mice to morphine disturbance. RL scores are calculated by determining the change (loss or gain) in ASV richness from a starting reference state (pre-disturbance) that occurs during a disturbance and subsequently relates this change or difference in richness to the eventual difference in richness between the reference and post-disturbance (recovered) states. In effect, the RL score attempts to capture the degree to which the microbiota returns to baseline richness after a change in richness in response to disturbance (see methods). Values range from “1” (complete recovery, reference diversity = recovered diversity) to “–1” (no change occurred during disturbance, reference diversity = disturbed diversity) (Supplemental Figure 5, Figure 3C). Because microbiota normally display day-to-day variability, three time points pre-paradigm where no systematic disturbance occurred (see Figure 3A) were used to determine the range of RL values that reflect normal variability (pink bar and points, Figure 3C) and establish the threshold for variation reflective of disturbance (black dashed lines, Figure 3C).

When the resiliency of microbiota diversity was determined from individual mice as they transitioned from the reference (“baseline”) state through morphine disturbance (“disturbed”) to post-disturbance (“recovered”) using the RL index, compulsive mouse microbiota averaged a positive RL value exceeding the threshold for day-to-day variability (p-value 0.043, Figure 3C, red bar), whereas non-compulsive mouse microbiota averaged a negative value that also exceeded the threshold for day-to-day variability (Figure 3C, grey). Though on its surface the positive RL value of the microbiota of compulsive mice could imply greater resilience (a partial return to starting diversity), and negative values suggest non-compulsive mice continued to experience more change post-disturbance consistent with a destabilized microbiota, the prior observed convergent loss of within-mouse microbiota variability (Figure 3A & B) and subsequent relatively greater resistance of the microbiota of compulsive mice to change post-morphine tempers this interpretation. Rather, non-compulsive mice appear to be in a state of transition from the convergent morphine state toward increased within- and/or between-mouse variability after removal of morphine. In contrast, the alternative state of the microbiota of compulsive mice was less variable both within and between mice post-morphine, reinforcing that the alternative state of the microbiota of compulsive mice may be comparatively stable.

We next evaluated the response of the post-morphine microbiota to disturbance and resiliency to recover from pulse morphine during the reinstatement trial that all mice experienced (Figure 3C). The ASV richness of disturbed and post-disturbed (“recovered”) microbiota of non-compulsive mice show a similar magnitude of change or divergence from the reference pre-disturbed state (population average ASV richness during extinction) in response to the pulse of morphine that was consistent with normal day-to-day variation (Figure 3C, grey bars, black dashed lines). In contrast, pulse morphine during the single reinstatement trial did not change the ASV richness of microbiota from compulsive mice, suggesting that the post-morphine microbiota in compulsive mice was resistant to pulse morphine disturbance as if it is adapted to morphine and displayed a greater magnitude of change in richness post-paradigm as indicated by the negative RL value (Figure 3C, dark red bars, black dashed lines).

### Distinct morphine-driven changes in complexity of microbiota define the alternative stable states of compulsive and non-compulsive mice

To shed light on the temporal patterns of microbiota diversity in responses to and recovery from morphine disturbance (Figure 3), we next examined whether genera-level microbiota population structures differed between compulsive and non-compulsive mice using networks analysis, which provides a perspective on the ecological complexity of the community. Coordinated abundance patterns of genera, which imply connections in the community by which information flows, were together organized as a network of interacting genera (Figure 4, genera are labeled with ASV number, ASV key is in Supplemental Table 3). Lines connecting genera (edges) are colored by whether the correlation of abundance is inverse (opposite association patterns; red) or direct (co-occurrence mirrors one another; green) and increasing weight of the line (density) represents higher frequency of detection of this connection among microbiota samples used to construct the network and thereby suggests stronger connections. Groups of genera that are most interconnected with each other signifying synchronization of abundances, are clustered in space and share color to form a module and together represent sub-communities corresponding to potential functional groups. The relative size of the colored circle directly reflects degree of Eigenvector centrality of the ASV, which is a measure of its importance to the community network structure, and the general position of the modules in the network also reflects centrality and influence within the larger network. Finally, genera whose abundances are synchronized with the greatest number of other genera in its associated module, and whose abundance is also sometimes synchronized with genera in other modules in the network, are identified as hubs (bold text) in that they act as a bridge for communication with other modules that form their own distinct functional groups. Ecologically, hubs can be seen as keystone genera that likely shape community structure and organization. The organization and interconnectedness of modules (modularity), the strength of connections, and degree of centrality or tightness of grouping can provide insight into robustness and presumed resiliency of the community as a whole. Though the analysis uses the 70 most abundant ASVs in the combined dataset of the two conditions to be compared, only genera that are connected to (i.e. abundance significantly correlated with) at least one other genera are included in the visualized network, where ASVs that are not connected in one of the two networks are colored grey.

**Figure 4:**
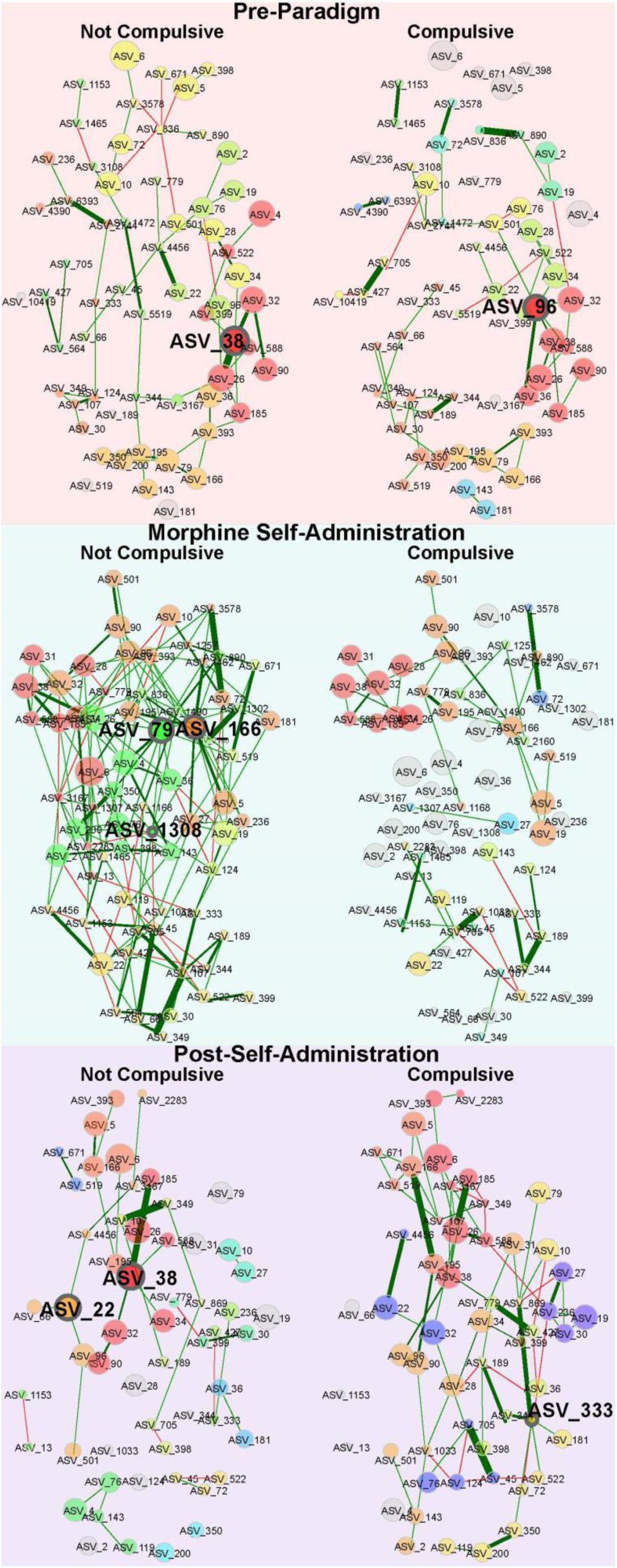
The microbiota networks of non-compulsive mice displayed greater connectivity and continuity during and after morphine whereas the networks of compulsive mice were fragmented and profoundly altered by morphine. Comparison of microbiota co-occurrence networks between compulsive and non-compulsive mice pre-paradigm (left panel), during morphine self-administration (middle panel), and after prolonged morphine during extinction, reinstatement, and post-paradigm (post-self-administration) (right panel) using the SPRING method in NetCoMi^59^. Edges are colored by sign (positive = green; negative = red), representing estimated associations between community members which are represented by colored circles with ASV listed and that are differentially colored by modules. Hubs are identified with bolded text of the ASV. Eigenvector centrality was used for scaling sizes of the circles representing genera (determined using greedy modularity optimization) where a shared color circle means the ASVs are highly connected to one another but have a small number of connections outside their group and assigned to the same module. Modules have the same color in both networks during pre-paradigm, during self-administration, and post-self-administration if they share at least two genera. Only genera that are connected in either compulsive and non-compulsive groups are included in the network, where grey circles indicate the ASV is not connected in one of the two parallel networks. Corresponding genera names detailed in Supplemental Table 3.

Side by side comparison of pre-paradigm networks indicate that the microbiota network properties of non-compulsive and compulsive mice did not differ (p-values for multiple global network properties detailed in the “whole network” subsection, Supplemental Table 3A) and shared highly similar network structures with many overlapping modules, which group some but not all of the same ASVs, (Figure 4 pre-paradigm). Interestingly, each pre-paradigm network had only one hub: *Oscillospira* (ASV_38) in non-compulsive mice, and in the same module of compulsive mice *Coprococcus* (ASV_96). In contrast, the networks of compulsive and non-compulsive mice diverged drastically during the self-administration phase in the number and density of edges connecting ASVs, the organization of ASVs to modules, the interconnectedness of modules, and the number of ASVs that served as hubs (bold text) (middle panel, Figure 4). The microbiota network of non-compulsive mice organized into five colored modules which displayed many within- and between-module connections representing a fully interconnected network, whereas the network of compulsive mice exhibited fewer and sparser connections where many modules and individual ASVs that were connected in non-compulsive mice were not connected to any others, representing a significant difference in overall network connectivity (p-value = 0.01982, Supplemental Table 3; Figure 4, middle panel).

The most central taxa in the network of non-compulsive mice during morphine self-administration included many taxa involved in digestion and gut health including *Ruminococcaceae* (ASV_79), *Bacteroides* (ASV_4), *Muribaculaceae* (ASV_6), *Christensenellaceae* (ASV_393), *Blautia* (ASV_398) and *Clostridium* (ASV_3167)^80-86^ (Supplemental Table 3). Some other central genera in non-compulsive mice are associated with morphine dysbiosis in mice, such as *Prevotella* (ASV_45), *Ralstonia* (ASV_1308), *Sphingomonas* (ASV_1490), and *Sutterella* (ASV_76, specifically identified as *Parasutterella excrementihominis*)^7, 15, 24^ (Supplemental Table 3). Except for *Prevotella* (ASV_45), these central genera were not connected to any other genera in compulsive mice during the self-administration phase (gray circles, middle panel, Figure 4; Supplemental Table 3). Moreover, only the microbiota network of non-compulsive mice contained hubs, including *Ruminococcaceae* (ASV_79), *Adlercreutzia* (ASV_166), and *Ralstonia* (ASV_1308). Curiously, the pre-paradigm networks grouped *Ruminococcaceae* (ASV_79) and *Adlercreutzia* (ASV_166) within the same peripheral module in both compulsive and non-compulsive mice, but in the morphine-disturbed network of non-compulsive mice, these both became central hubs of distinct novel modules, as denoted by different colors.

Comparison of the microbiota network structures of compulsive and non-compulsive mice following the removal of morphine also revealed divergent network patterns, most notably in edge density signifying strength of co-occurrence relationships as well as overall connectivity, and connectedness of core genera and modularity (Figure 4 and Supplemental Table 3). The microbiota networks of compulsive mice had a tight core with significantly higher edge density (p-value = 0.040, Supplemental Table 3, Figure 4) representing strong connections between the most central genera (centrality metrics, Supplemental Table 3C), consistent with their dominance over network structure. Together these data suggest a high degree of network robustness. However, despite their overall stronger natural connectivity through the influence of this core, peripheral functional groups were less connected (modularity, p=0.030) in the networks of compulsive mice compared to the less tightly linked network of non-compulsive mice, and this lower modularity could stymie crosstalk between different sub-communities. This pattern of central interconnected prominence marks a reversal from the patterns evident from comparisons between networks of non-compulsive and compulsive mice during morphine self-administration, when non-compulsive mice had the more central and connected core (see centrality measures for the most central nodes, Supplemental Table 3B & C). This suggests that the microbiota of compulsive mice may have undergone a reorganization or strengthening of core interactions in response to morphine removal, potentially reflecting a lingering or compensatory network response to the morphine experience, one that persists even after weeks of abstinence. Although the overall network connectedness was higher in compulsive mice post-self-administration, the networks in non-compulsive mice showed signs of potential rebound, with greater modularity of microbial sub-communities, potentially allowing adaptive responses across functional groups. For instance, microbiota networks in non-compulsive mice regained *Oscillospira* (ASV_38) as a hub, which was notably the only pre-paradigm hub in the microbiota of non-compulsive mice and these networks also maintained many of the pre-paradigm modules through and after morphine self-administration, indicating continuity in sub-community organization. The microbiota networks in non-compulsive mice gained *Rikenellaceae* (ASV_22) as a new hub connecting its module to the central module containing the hub *Oscillospira* (ASV_38). The most central genera forming new connections in compulsive mice included genera considered beneficial and include *AF12* (ASV_344), *RF39* (ASV_28), *Clostridiales* (ASV_26), *Odoribacter* (ASV_189), and *Bifidobacterium* (ASV_27)^7, 15, 82, 87-94^. However, several new modules formed that were dominated by potential pathobionts (*Desulfovibrio* (ASV_124*), Streptococcus* (ASV_869), and *Alphaproteobacteria* (ASV_333), *F16* (ASV_1033), and *Coriobacteriaceae* (ASV_236))^15, 20, 95-102^ (Figure 4, right panel; Supplemental Table 3), suggesting these networks are dysbiotic. Even as the microbiota of non-compulsive mice regained some of its pre-morphine organization, they too formed novel modules post-morphine (Figure 4, purple, blue and aqua) connecting *Turicibacter* (ASV_10) and *Bifidobacterium* (ASV_27) which were previously connected only in the pre-morphine networks of compulsive mice.

The notable differences in the formation of modules (Figure 4) suggest potential different changes in functional diversity in response to morphine disturbance in compulsive and non-compulsive mice. To assess this, functional profiles of predicted metabolic pathways were generated using the PICRUSt 2 pipeline^60^ and differences in the abundance of predicted metabolic pathways between non-compulsive and compulsive mice were identified using linear modeling (LinDA)^66, 103^ (Supplemental Figure 6). No differences in metabolic capacity were identified pre-paradigm, which aligns with the lack of significant differences in microbiota structure between non-compulsive and compulsive mice (Figure 4). However, differences emerged during morphine self-administration, where metabolic pathways involved in short chain fatty-acid (SCFA) biosynthesis, the generation of precursor metabolites and energy, nucleotide biosynthesis, carbohydrate metabolism, cofactor biosynthesis, nitrogen metabolism, polysaccharide degradation, heme biosynthesis, fatty acid biosynthesis, and aromatic compound degradation were significantly enriched in non-compulsive mice (Supplemental Figure 6), suggesting a functionally more diverse microbiota than in compulsive mice. In contrast, compulsive mice were only enriched in metabolic pathways involved in amino acid biosynthesis, and one pathway in nucleotide biosynthesis and carbohydrate metabolism (Supplemental Figure 6), suggesting a greater loss of functional diversity in response to morphine disturbance compared to non-compulsive mice. Following the removal of morphine, non-compulsive mice better maintained broad functional diversity as evidenced by the enrichment of predicted functional pathways associated with SCFA biosynthesis, the generation of precursor metabolites and energy, carbohydrate metabolism, heme biosynthesis, and nitrogen metabolism, as compared to compulsive mice (Supplemental Figure 6).

### Compulsive mice exhibited a relatively greater decline in beneficial genera and increase in potential pathobionts

Since the microbiota of non-compulsive and compulsive mice responded differently to morphine disturbance (Figure 3 & 4), we next identified microbiota community members whose abundance significantly correlated with behavioral differences, but limited comparisons to mice with the most extreme phenotypes (composite compulsivity scores above the population average + 2 IQD indicated the most compulsive mice and composite scores below the population average – 2 IQD indicated the least compulsive mice, dashed lines in Figure 5 inset) in order to reduce potential noise from overlap in microbiota and behaviors among the intermediately scoring mice that could obscure relationships (Figure 1E). Differentially abundant genera were identified using a beta-binomial regression model^67^ (Supplemental Table 4, see methods for details) and then from among the differentially abundant genera, biomarkers, or genera that were significantly correlated with and most explained the difference between the mice with the highest and lowest compulsivity scores were identified. Overall, relatively few biomarkers distinguished the most compulsive from least compulsive mice (Figure 5, see Supplemental Table 4 for details and statistics). Whereas no distinguishing biomarkers were identified pre-paradigm, a few biomarkers, more in the most compulsive mice, emerged early during morphine self-administration (weeks one to 12, grey and red circles, Figure 5). Most biomarkers emerged during late morphine self-administration (weeks eight to18) indicating significant divergence of the microbiota of the most and least compulsive mice. More of these were associated with the least compulsive mice and included well-known probiotics like *Bifidobacterium* (ASV_27) and *Parabacteroides* (ASV_66)^104^, and others with potential benefits for promoting gut homeostasis, such as *Oscillospira* (ASV_38), *Ruminococcus* (ASV_90), *Adlercreutzia* (ASV_166), and *Coprococcus* (ASV_96)^94, 105-108^ (Figure 5, grey circles). Several of these were also identified as global biomarkers of the least compulsive group due to their strong association even where microbiota from all stages of the paradigm were combined (grey circles next to italicized labels, Figure 5). Conversely, the three biomarkers that emerged late during self-administration in the most compulsive group are all identified as potential pathobionts in mice and included *Prevotella* (ASV_30), *Sutterella* (ASV_76), and *Coriobacteriaceae* (ASV_236)^7, 10, 15, 24^ (Figure 5, red circles). These three genera were also identified as global biomarkers as was *Muribaculaceae* (ASV_6) (Figure 5, red circles next to italicized names). Notably, *Muribaculaceae* (ASV_6) was one of the only biomarkers of the most compulsive mice that is not recognizable as a potential pathobiont but a common commensal. Although *Akkermansia muciniphila*, which was also a biomarker of compulsive microbiota, is a probiotic species, it has been associated with some disease states^109^. Following morphine self-administration during extinction, reinstatement, and post-paradigm, only two genera, both considered beneficial taxa were biomarkers of the least compulsive mice (grey circles, Figure 5). In contrast, *Clostridium* (ASV_143) emerged as a new biomarker of the most compulsive mice during extinction, while *Prevotella* (ASV_30) remained elevated in the most compulsive mice and was a biomarker during the extinction and reinstatement phases and post-paradigm (red circles, Figure 5).

**Figure 5:**
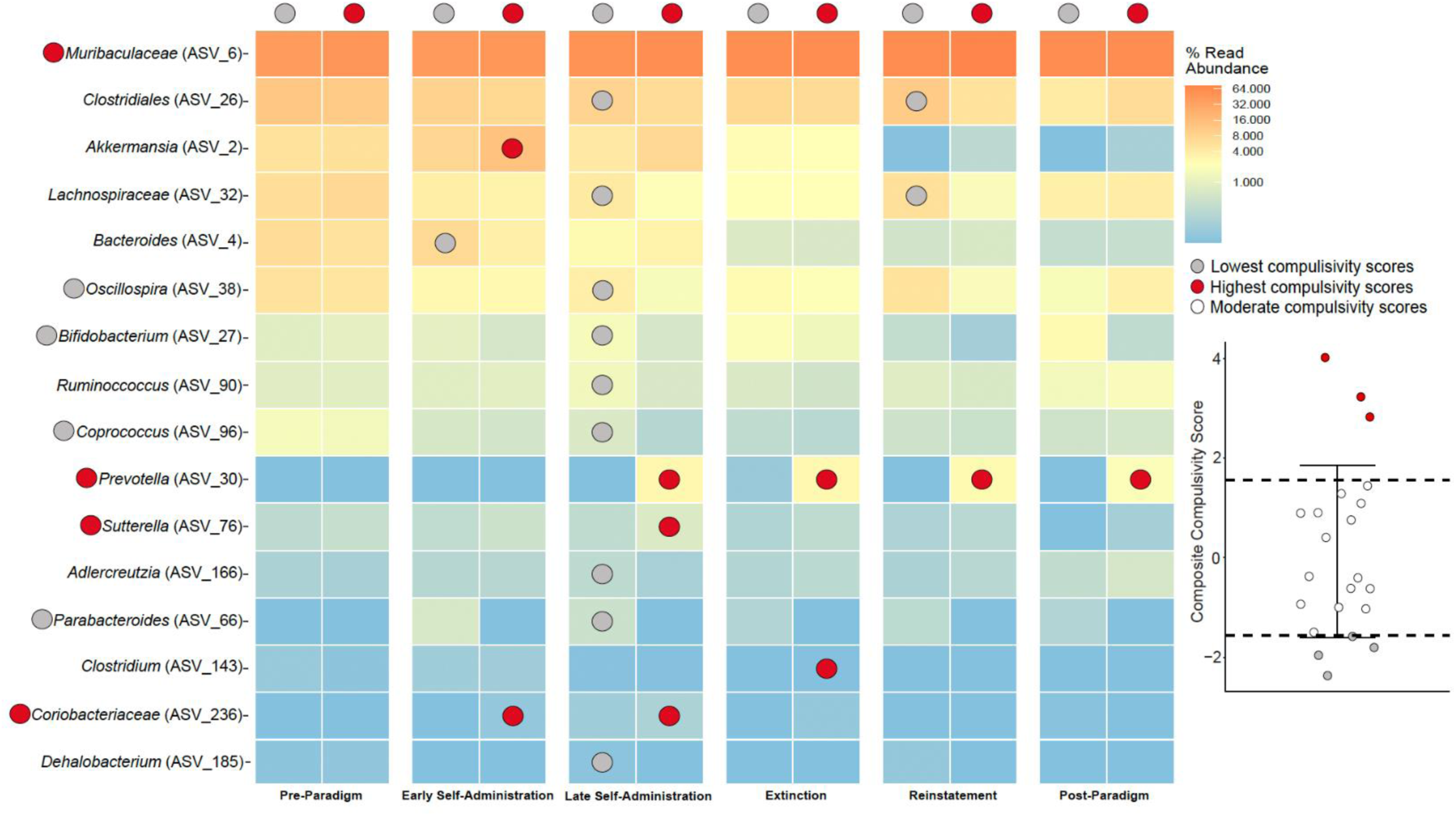
Microbiota associations with extreme compulsivity scoring mice. Biomarkers of the most compulsive (n = 3 mice, red circles) and least compulsive (n= 4 mice, grey circles) --see inset-- were identified from among the community members whose abundance was explained by categorical assignment (as identified using corncob regression models)^67^, followed by a linear discrimination analysis (LDA) of effect size (LEfSe)^68, 69^ representing genera that most likely explain differences between the microbiota in each category. Mice with scores above the population average + 2 IQD (red circles, inset distribution) or with scores below the population average - 2 IQD (grey circles, inset distribution) were used in this analysis, whereas mice with moderate scores (white circles, inset distribution) were not used for biomarker identification. Community members are labeled at the genus level or at the lowest classification available when genus assignment was not available. * by the genera name were identified as biomarkers when microbiota from all phases of the paradigm were combined. See Supplemental Table 4 for details.

Next, we examined whether there were unique memberships, or indicator genera, associated with categorical assignment of mice as compulsive or non-compulsive^71^. Only 10 indicator genera were identified, of which most were associated with compulsive mice (80%, Supplemental Table 5, Supplemental Figure 7). Surprisingly, among the eight genera that were indicators of compulsive mice, most (75%) were identified pre-paradigm or early during morphine self-administration. Specifically, *Sporobacter* (ASV_2283), *Candidatus arthromitus* (ASV_427), and *Curtobacterium* (ASV_1462), were differentially associated with compulsive mice pre-paradigm, and *Peptococcaceae* (ASV_705), was differentially associated with compulsive mice both pre-paradigm and early during morphine self-administration. None of these were exclusively associated with compulsive mice as each subsequently increased to the detection limit in non-compulsive mice after prolonged morphine exposure. Conversely, indicators of compulsivity during extinction and reinstatement were identified as such because their levels fell below detection in non-compulsive mice, alluding to how the communities were shaped differently by morphine, but also suggesting some subtle differences even prior to morphine.

The longitudinal paradigm integrates multiple behavioral measures related to effort to obtain drug (Figure 1), but analyses thus far have used vernacular categorical assignment of mice as justified statistically (Figure 1C & E and Figure 5) to provide more robust statistical support for inferences. But this categorical assignment fails to capture that mice with intermediate phenotypes and categorized as non-compulsive may have undergone intermediate changes that could help clarify which community members are most associated with behavioral shifts. To address this limitation, we next identified how relative abundances of genera correlated with individual compulsivity scores as a continuous variable (Figure 6). Correlations of individual genera abundance with composite compulsivity scores were determined, their significance evaluated (see methods), and these results summarized with directionality of correlation plotted horizontally from center, effect size indicated by size of circle, and statistical significance above the default threshold for significant correlation (inverse log q = 0.6) scaled vertically (Figure 6, Supplemental Figure 8). These analyses allowed us to assess microbiota association patterns identified by categorization that align with those that are based on continuous measures where low scores indicate a mouse shows generally less lever pressing, quick extinction and low reinstatement, and higher scores indicate a mouse shows generally more lever pressing, longer extinction and robust reinstatement (Figure 6, Supplemental Figure 4).

**Figure 6:**
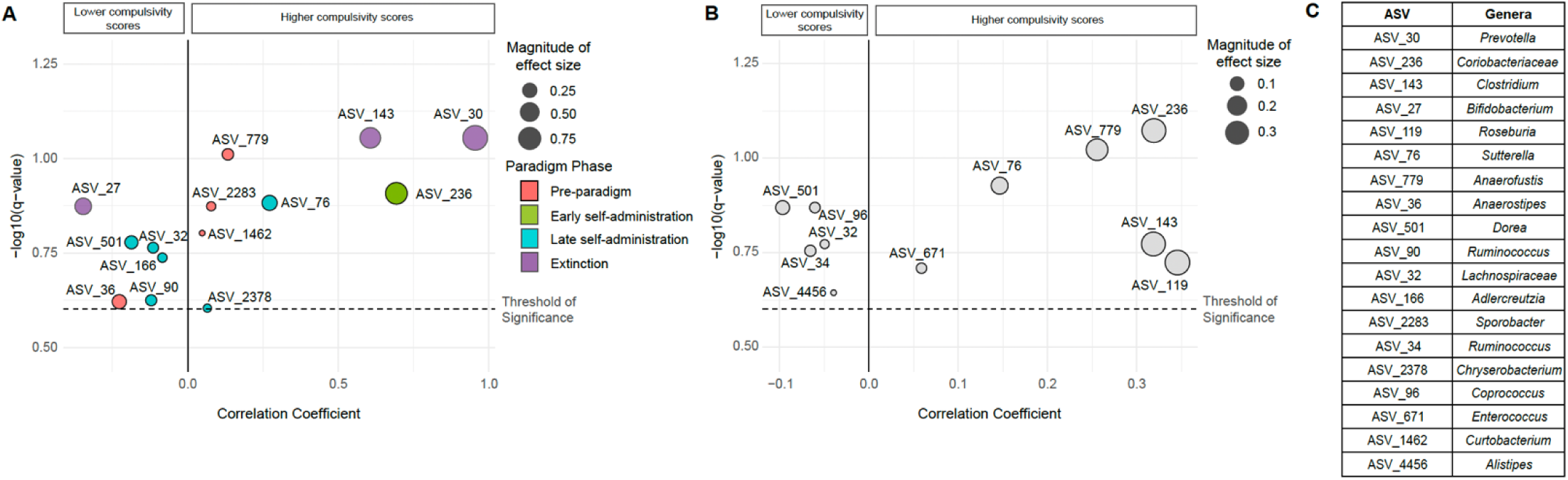
Gut microbiota genera were more strongly correlated with higher compulsivity scores than lower compulsivity scores. A) Associations of center-log ratio (CLR) transformed relative abundances of individual genera (labeled as ASV) with composite compulsivity score using microbiota from a given stage or phase where circles representing effect size of genera are colored by stage or phase of the paradigm including pre-paradigm (orange circles), early self-administration (green circles), late self-administration (blue circles), and extinction (purple circles) were assessed using multivariable association testing with linear models (MaAsLin 2)^72^. Negative correlation coefficient values indicate inverse correlations with composite compulsivity scores, and positive correlation coefficient values represent direct correlations with composite compulsivity scores distributed horizontally. The size of individual colored circles correspond to the magnitude of effect size, or the strength of the correlation. B) Associations of CLR transformed relative abundances of individual genera (labeled as ASVs) from all phases combined (light grey circles) with composite compulsivity scores were assessed using multivariable association testing with linear models (MaAsLin 2)^72^. Only genera identified as significantly correlated based on the default threshold (q-value of 0.25, noted as the inverse log q = 0.6) as indicated by the dashed black line, are presented. Negative correlation coefficient values indicate inverse correlations with composite compulsivity scores, and positive correlation coefficient values represent direct correlations with composite compulsivity scores distributed horizontally. The size of individual grey circles corresponds to the magnitude of effect size, or the strength of the correlation. C) Genera names for ASV labels from A and B are organized in descending order of magnitude of effect size. See Supplemental Figure 8 for correlations, including those below the significance threshold.

Genera that correlated with higher compulsivity scores had larger effect sizes indicating stronger correlations. These include *Coriobacteriaceae* (ASV_236), a potential pathobiont^110^ identified during self-administration (green circle Figure 6A), *Sutterella* (ASV_76), a biomarker of morphine dysbiosis^24^, during late self-administration (blue circle, Figure 6A), and the strongest correlating genera *Prevotella* (ASV_30) and *Clostridium* (ASV_143) during extinction (purple circles, Figure 6A). Although a similar number of genera correlated with lower and higher composite compulsivity scores when all microbiota were analyzed as a combined dataset (five and six respectively) the pattern of strongest correlations with compulsivity was also conserved (Figure 6B). *Bifidobacterium* (ASV_27) a known probiotic genera showed the strongest correlation with low compulsivity scores and was also identified during the extinction phase (purple circle, Figure 6A), and many other known beneficial genera also correlated with low compulsivity scores (e.g. *Adlercreutzia* (ASV_166), *Bifidobacterium* (ASV_27), *Ruminococcus* (ASV_90), and *Ruminococcus* (ASV_34)). No significant correlations between the abundance of genera with the compulsivity score were apparent during reinstatement or post-paradigm, but a few genera identified from pre-paradigm microbiota directly but relatively weakly correlated with higher compulsivity scores, and two of these were also indicator genera (*Sporobacter* (ASV_2283) and *Curtobacterium* (ASV 1462)). Only a subset of genera previously identified using categorical assignment of compulsivity also showed significant correlations with compulsivity score as a continuous variable, though some new genera were also revealed in this approach and in all phases of the paradigm (Table 1). These findings reinforce that key distinguishing microbiota signatures include an increase in potential pathobionts during and post-morphine disturbance is apparent among mice as they develop compulsive behaviors, whereas higher abundance of potentially beneficial genera are associated with lower compulsivity.

**Table 1:**
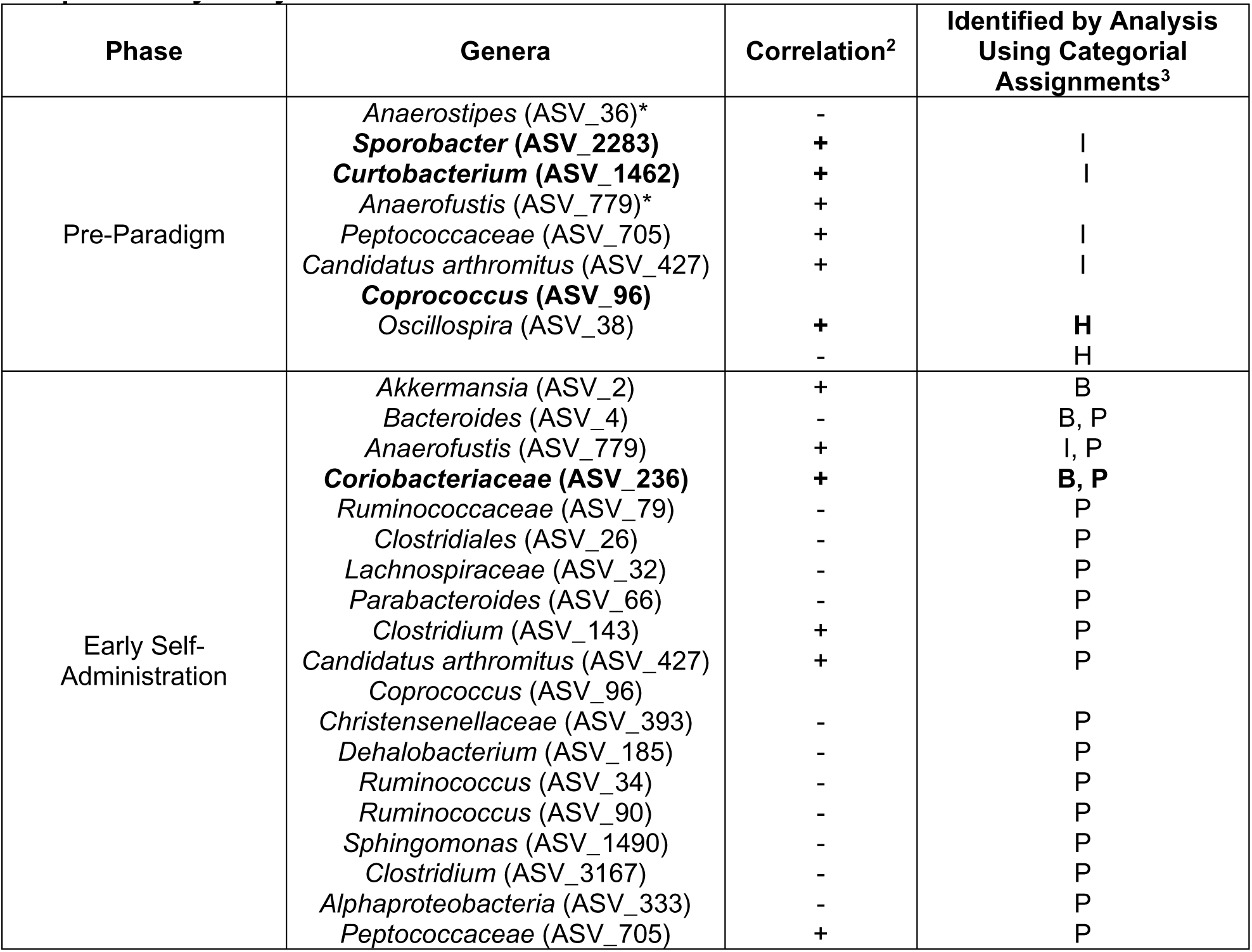
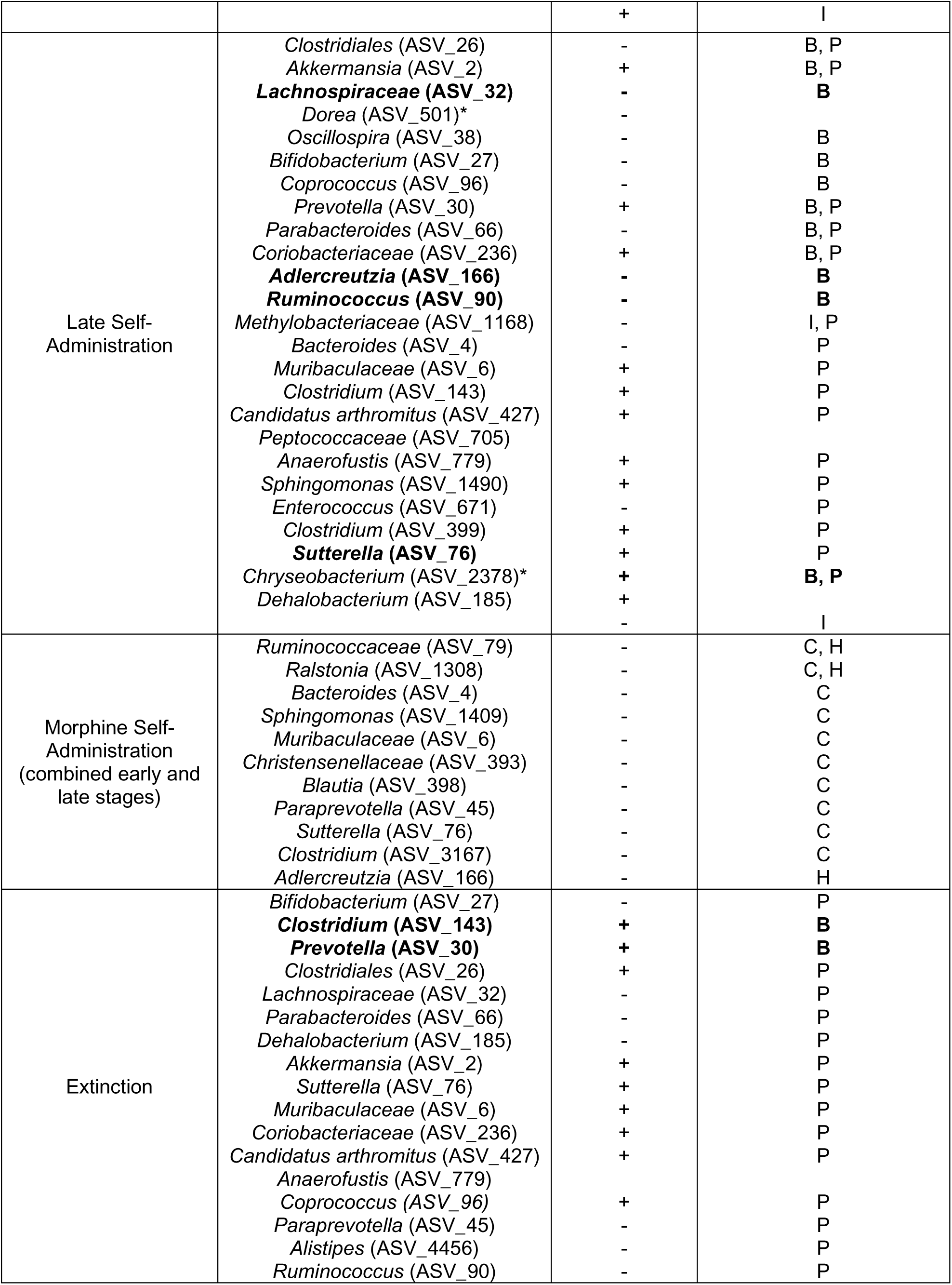
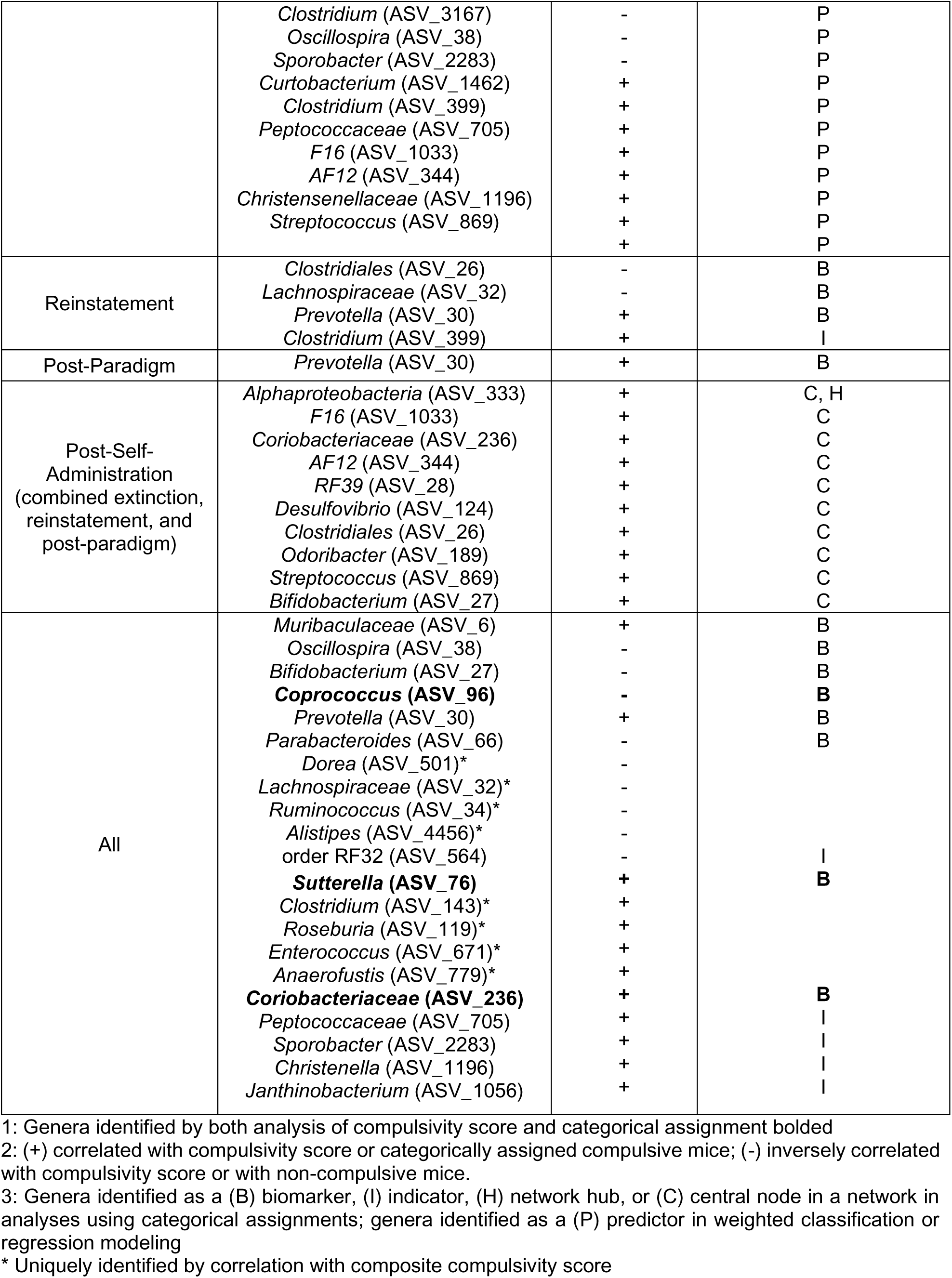
Summary of genera correlated with compulsivity score or identified by complimentary analyses^1^.

### Genera predictive of compulsivity formed stable associations in compulsive mice and dynamic associations in non-compulsive

Our prior analyses using either categorical assignment or compulsivity score as a continuous variable revealed microbiota signatures that distinguished compulsive mice from non-compulsive mice that suggest a temporal pattern of divergence during morphine self-administration but also post-morphine. To further examine whether the distinctive temporal microbiota association patterns could predict the final behavioral outcome, thereby providing additional support that these differences were linked to behavior, we applied supervised machine learning to iteratively build models to distinguish between microbiota of mice not only when they were categorized as compulsive or non-compulsive (weighted classification) but also to predict the degree of compulsivity (regression) based on the random forest algorithm^74^. Composite compulsivity scores and associated microbiota compositional data were split into training sets for model generation and testing sets for validation (see methods for details). We assessed the reliability and accuracy of each model in distinguishing compulsive from non-compulsive mice (categorical/weighted classification) or for predicting the individual numeric compulsivity score (continuous/regression) using multiple performance metrics, including those that assess accuracy for categorical assignment, and that evaluate how accurately regression models predicts, or explains the variance in compulsivity score in comparison to a mean-prediction model (Supplemental Figure 9). Neither model performed sufficiently when using microbiota from pre-paradigm, the reinstatement phase or post-paradigm (i.e. genera abundances were not predictive during these phases, Supplemental Figure 9). In contrast, the weighted classification and regression models performed optimally in predicting compulsivity category and score with microbiota collected both early and late during the morphine self-administration phase and the extinction phase (i.e., genera abundances were predictive during these phases, Supplemental Figure 9). Sample size did not explain the observed differences in model performance; rather, stronger and more consistent microbiota shifts during morphine self-administration and extinction likely account for better model performance during those phases. We then identified the genera that contributed most to model performance (Supplemental Figures 10, 11, Figure 7). Because the random forest algorithm used in both models does not provide directionality in the relationship between community members and outcomes (e.g., whether a community member correlates with or against compulsivity score), we cross-referenced genera identified in the models with genera from our previous analyses (Figures 4, 5 and 6, Table 1, Supplemental Tables 4, 5) providing directionality and interpretability to predictive genera.

**Figure 7:**
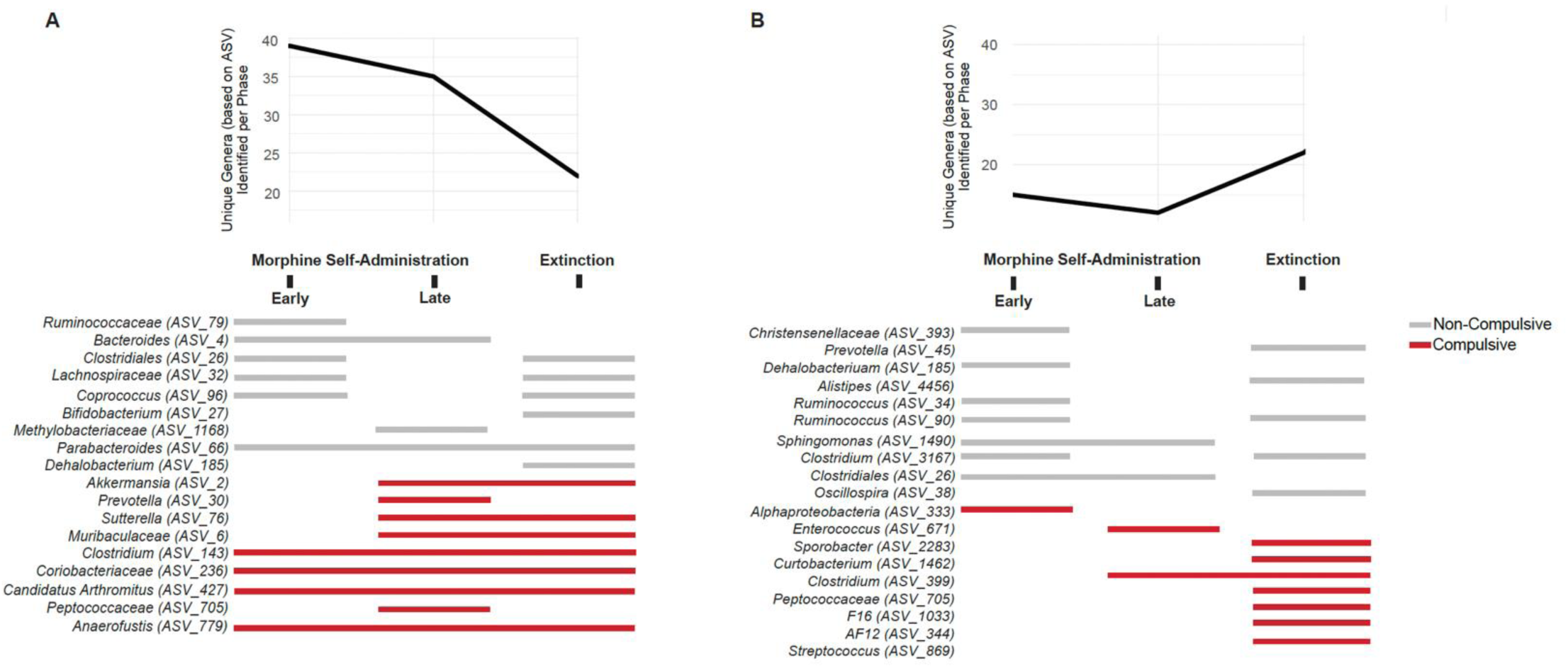
Predictive genera formed dynamic associations in non-compulsive mice, contrasted with more stable and persistent associations in compulsive mice. A) The total number of unique genera identified per phase during weighted classification modeling (using categorical assignments) using permutation testing in mikropml^74^ (top), and identification of genera that are most important for the generation of random forest prediction models to classify mice as either compulsive (red) or non-compulsive (grey) at specific phases of the paradigm using mikropml^74^ and that were also identified by previous analyses (bottom panel) using compulsivity as a categorical variable (i.e., biomarkers and network hubs) to determine whether a predictive community member most likely originates from a microbiota of a compulsive (red) or non-compulsive (grey) mouse. See Supplemental Figure 10 for the complete list of genera important for model performance. B) The number of unique genera identified per phase during regression modeling (using composite compulsivity scores) using permutation testing in mikropml^74^ (top) and identification of genera that are most important for the generation of random forest prediction models to associate with the degree of compulsivity at specific phases of the paradigm using mikropml^74^ that were identified by other analyses using compulsivity as a continuous variable (i.e., multivariate association with linear models) to associate whether a predictive genus most likely originates from the microbiota of a mouse with a lower or higher compulsivity score (bottom panel). See Supplemental Figure 11 for the complete list of genera important for model performance.

In categorical assignment models, several of the cross-referenced genera associated with the microbiota of compulsive mice during early morphine self-administration were already predictive of the eventual categorical assignment of these mice as compulsive and all of these genera remained predictive of compulsive mice through extinction suggesting early divergence in the microbiota could have precipitated decline but they persisted thereafter (red lines, Figure 7A). Several more genera emerged as predictive in the microbiota of compulsive mice during late morphine self-administration aligning with “disease” progression and most of these also persisted during extinction (red lines, Figure 7A). In contrast, most of the genera predictive of non-compulsive mice were only predictive during one phase with few spanning more than one phase and most were either associated with early morphine self-administration or extinction (grey lines, Figure 7A). The only predictive genera to span consecutive phases include *Bacteroides* (ASV_4) during early and late morphine self-administration, and *Parabacteroides* (ASV_66) during morphine self-administration and extinction. The general lack of predictive associations during late morphine self-administration suggests if beneficial genera were bolstering the microbiota of non-compulsive mice they were variable from each other.

In models predicting compulsivity score, relatively more cross-referenced genera (seven) inversely correlated with compulsivity score (lower compulsivity score) than directly correlated (one) during early self-administration which could reflect that decline of these genera were predictive of a path toward compulsive behaviors (Figure 7B). Some of these genera predictive of low compulsivity scores spanned into late self-administration and several new genera emerged as predictive during extinction. In contrast, more genera associated with extinction were predictive of high compulsivity scores, with only a few emerging during morphine self-administration (red lines, Figure 7B). Overall, visual exploration of the temporal distribution of genera contributing to these models suggest that genera predictive of the compulsive category and higher compulsivity scores formed stable associations and often remained predictive through or sometimes even emerged during extinction, whereas genera predictive of the non-compulsive category and lower compulsivity scores showed more dynamic predictive relationships as one might expect under conditions of equilibrium and reflective of functional redundancy. Additionally, the higher number of genera predictive of categorical assignment during early morphine self-administration (∼40) (Figure 7) compared to other analyses using categorical assignments (five) (Figure 5 and Supplemental Table 5) may also indicate dynamics of early morphine disturbance could be relevant.

## DISCUSSION

Expanding upon our prior study characterizing morphine-induced microbiota responses that were tied to--and protective of--the development of antinociceptive tolerance^15^, here we show that differences in microbiota composition correlate with behaviors that underlie OUD as a chronic and relapsing condition. Specifically, we modelled the progression from impulsive to compulsive drug-seeking using a three-phase murine behavioral paradigm^16^ that aligns with multiple DSM-5 criteria^4, 5^ to generate a continuous composite score reflecting time and effort to obtain drug and drug craving, and leveraged natural variation in microbiota and behavior^15, 16^ to examine correlations between the two variables. This revealed that despite morphine shifting the microbiota of all mice to an alternative stable state (Figure 2 & 3), the morphine-disturbed microbiota of mice categorized as non-compulsive or with lower compulsivity scores displayed relatively more dynamic flexibility reflected in enhanced within- and between-mouse variability of diversity and variable predictive associations (Figure 3 & 7), community complexity and interconnectedness (Figure 4), and broader functional diversity associated with beneficial genera (Supplemental figure 6) both during morphine press disturbance and in the post-morphine equilibrium state of microbiota. In contrast, morphine restructuring of the microbiota of mice categorized as compulsive and with higher compulsivity scores convergently produced a less functionally diverse microbiota dominated by potential pathobionts (Figure 4 and 5) and the association of these genera, especially during late self-administration and persisting though extinction, were strongly correlated with and predictive of compulsive behavior (Table 1, Figure 6 & 7). It was especially apparent that the compensatory community structure that remained post-morphine in these compulsive mice was robustly unyielding (Figure 3 & 4) and “stuck” in a dysbiotic state. These contrasting patterns in native microbiota responses to morphine point to the potential role of the gut microbiota in modulating opioid-induced neurobiological changes either for the better or the worse.

OUD is a complex syndrome characterized by loss of control over drug seeking and craving, and relapse during abstinence that can persist long after drug use has ceased (for review see^111^). In humans, OUD is diagnosed based on a series of 11 criteria most recently defined by the DSM-5^4^. Those positive for 2-3 of these criteria have a mild OUD, 4-5 a moderate OUD and 6 or more severe OUD. Importantly, individuals can present with any combination of symptoms, which means a patient with tolerance and physical dependence but none of the other criteria (2 symptoms) and a patient with craving and who takes more opioids than intended but no other criteria (also 2 symptoms) would both be diagnosed with a mild OUD. Hence, the presentation of the disorder varies greatly across the affected population and likely reflects differences not only in behavior but the actual neurobiological changes that led to the disorder. Modeling such a complex disorder in rodents is a challenging task. Indeed, some of the DSM-5 criteria cannot be modeled in rodents at all (taking more drug than *intended* for example)^4, 5^. While many rodent studies use reward or operant self-administration as a proxy for OUD, neither drug reward nor drug taking is a criteria of human OUD per the DSM-5^4^. Therefore, to best model OUD in rodents, it is critical to capture as many aspects of the disorder as possible. The model we present here is designed to both allow for voluntary consumption of large quantities of drug across many weeks—sufficient to produce both tolerance^15^ and dependence^16^ in most animals--and to interrogate several additional aspects of OUD, including drug seeking, drug craving, and relapse. As we do for human OUD, we then created a diagnostic tool for assessing severity of OUD-like symptoms in the mice that we termed their “compulsivity score”^16^. Critical to our studies here, in which we are assessing the role of the microbiota in OUD, this model preserved variability across the population in the severity of symptomology, akin to mild to moderate to severe OUD-like behavior. Most prior studies of opioids and the rodent microbiota have collapsed behavioral variability, primarily by administering the same dose of non-contingent morphine to each subject ^7, 10-12, 21, 25, 112^. Leveraging the variability in mouse behaviors through many weeks of voluntary opioid administration and the innate variability of the individual microbiota was instrumental in our ability to identify the changes in microbiota that co-occur with either protection from or the promotion of OUD-like behavior.

The microbiota of compulsive mice was profoundly disrupted by morphine exposure, with changes progressing to a highly persistent state with limited variability suggesting they lost some adaptive capacity (Figure 1A, Figure 5, 6 and 7). The persisting state was marked by depletion of beneficial genera and loss of their functional diversity, and concurrent dominance by potential pathobionts both of which are common signs of morphine exposure^6, 7, 9, 10, 15, 20, 21, 23, 24, 112-115^. Across analyses, including those revealing that indicator genera of compulsivity pre- and early morphine were typical commensals (Supplemental Table 5), and comparisons of starting composition that revealed few differences (Figure 2, 5, 6, 7) there was no indication that the microbiota of compulsive mice was predisposed to failure in response to morphine. Notably, even though the network properties suggest no difference between compulsive and non-compulsive mouse microbiota pre-morphine, curiously they were anchored by different hub genera: *Oscillospira* (ASV_38) in non-compulsive networks and *Coprococcus* (ASV_96) in compulsive networks (Figure 4, Supplemental Table 3). Even though both of these are commensals associated with promoting gut homeostasis and microbiota diversity^105, 106, 116, 117^, their role in shaping microbiota structure following morphine disturbance diverged, as *Oscillospira* re-emerged as a hub in the non-compulsive network post-self-administration whereas *Coprococcus,* a genera susceptible to morphine disturbance^15^ did not (Figure 4). Rather, the microbiota structure in compulsive mice became sparse (Figure 4), with grave consequences since other beneficial commensals that could have potentially promoted recovery were also depleted relative to their abundance in non-compulsive mice, indicating the microbiota community structure that formed without this hub did not counteract progressing dysbiosis (Figure 4, Figure 5). However, as a biomarker in non-compulsive mice during late self-administration (Figure 5), *Coprococcus* could contribute to structure as a beneficial commensal, but it was perhaps not as effective as a keystone genera. Two new modules comprised solely of known pathobionts emerged in the compulsive network post self-administration (purple and blue, Figure 4), and included *Alistipes* (ASV_4456), *Paraprevotella* (ASV_45), *Desulfovibrio* (ASV_124), *Sutterella* (ASV_76), *Coriobacteriaceae* (ASV_236), and *Prevotella* (ASV_30)^7, 10, 15, 24, 95, 110, 118-121^. Abundance of some of these--especially during late self-administration and post-morphine--correlated strongly with composite compulsivity scores (Figure 6A), along with *Clostridium* (ASV_143) which from this association, and that it was also a biomarker of the microbiota of the most compulsive mice during extinction, raises suspicions of its potential role^82, 122^. Additionally, the networks of compulsive mice formed new sub-communities centered around *Alphaproteobacteria* (ASV_333), a potential pathobiont^96^, which emerged as a keystone genus in the post-self-administration network (Figure 4). This stronger shift towards pathobiont dominance indicate persistent dysbiosis^30^. Further assessment of genera whose abundance is strongly correlated with compulsivity score, with a large effect size especially during late self-administration and post-morphine including Sutterella (ASV_76) (specifically *Parasutterella excrementihominis*), *Coriobacteriaceae* (ASV*_*236), and *Prevotella* (ASV_30) is certainly warranted as their association with persisting behaviors could reflect ongoing modulation of behavior by the dysbiotic microbiota.

In contrast, even as non-compulsive mice experienced morphine disturbance that changed microbiota diversity and composition (Figure 2, 3 and 5), their microbiota nonetheless were characterized by partial “recovery” of normal variability (Figure 3), more dynamic community interactions and associations during press morphine with more variable predictive capacity (Figure 4, 5, 6 and 7) but at the same time with comparatively more continuity in community structure through and after press morphine disturbance (Figure 4) indicating an adaptive microbiota better able to respond to recurring morphine disturbances. Non-compulsive mice harbored relatively higher abundances of many beneficial genera (e.g. *Bifidobacterium* (ASV_27), *Bacteroides* (ASV_4), and *Parabacteroides* (ASV_66)) that support gut-brain axis signaling and regulation of bile acid metabolism to maintain microbiota homeostasis, gut barrier integrity, and immune function^112-116^. Via their production of β-glucuronidase, an enzyme that deconjugates morphine metabolites for reabsorption, these community members may also modify morphine bioavailability^10, 20, 89^. During morphine disturbance the interconnectedness of the networks of non-compulsive mice was enhanced and organized around three keystone species (hubs) (*Ruminococcaceae* (ASV_79), *Adlercreutzia* (ASV_166), and *Ralstonia* (ASV_1308)). *Ruminococcaceae* and *Adlercreutzia* were additionally predictors of low compulsivity score and categorical classification as non-compulsive during early morphine self-administration suggesting a pivotal role for counteracting morphine’s destabilizing effects on community functional organization (Figure 7). Whereas these associations readily align with the potential that these community members effectively blocked more severe pathobiont overtake by exclusion, and that the pathobionts are at the root of microbiota association with compulsivity, complementary possibilities exist. For example, all three hub genera, and two other module-associated genera (*Bifidobacterium* (ASV_27) and *Ruminococcus* (ASV_90) (Figure 4)) that were also biomarkers associated with the least compulsive mice (Figure 5) and lower compulsivity scores (Figure 7) share a common metabolic property : their known capacity for bioactivation of soy isoflavones to produce the phytoestrogen equol^123, 124^. As a ligand of estrogen receptors throughout the body including in the brain in both male and female mice^125-130^ equol could potentially enhance estrogen receptor signaling to decrease severity of compulsive behaviors^131-133^ as has been documented with estrogen supplementation in male mice^131^. Whereas equol binding may reduce opioid reward and drug-seeking behavior, estrogen signaling and its effects on compulsivity are complex and sex-lined and estrogen has also been implicated in increased compulsivity in female mice^132, 134^. As intriguing as this potential link is, the mechanisms and interventions for modifying behaviors would need further experimental evaluation, though are quite feasible ^131^. It remains unclear how behavioral changes, if modified via equol, are connected to the overall lower degree of dysbiosis that is apparent in the non-compulsive mice who benefitted from these keystone species (Figure 4). At a minimum, this highlights how multi-faceted the role of the microbiota could be in OUD-where the same community members could both directly stabilize the community structure to curtail emergence of pathobiont centered functional communities (Figure 4, 5, & 6) and simultaneously influence MOR signaling in the host, which in turn could further impact community structure by localized changes in the gastrointestinal environment. Regardless, these associations suggest the potential of multiple intersecting roles of the microbiota especially of beneficial community members, in countering the adverse effects of morphine that associated with compulsivity.

The rise of pathobionts in response to morphine disturbance is most often attributed as the cause for microbiota exacerbation of opioid-induced behavioral changes for which they correlate-a concept that has gained traction due to studies that demonstrate amelioration of these effects in germ-free mice^7, 9^ and with antibiotic depletion of the microbiota^6, 7, 9, 12^. Indeed, the association remains at the forefront among the mice categorized as compulsive in our study. However, the response of microbial communities to enhance networks of communication during morphine disturbance and the notable preservation of beneficial genera cannot be ignored as important and reinforces that loss of supportive microbiota functions are also likely to contribute to behavioral changes. As previously shown, both probiotics^7^, and dietetic butyrate supplementation^15^ to replace metabolic functions lost during morphine depletion of beneficial genera counteract morphine-induced antinociceptive tolerance. However, if loss of beneficial genera played a role in compulsive behaviors, the uncoupling of compulsivity and tolerance (Supplemental Figure 4C) suggests microbiota responses that contribute to compulsivity could reflect distinctive processes from those that underlie tolerance. This is not entirely surprising since tolerance and linked behavioral changes such as dependence^9-12^ and hyperalgesia^21, 28^, can resolve rapidly once drug is removed, but, as we model here, compulsive behaviors persist long after drug is removed and contribute to the risk of relapse. Even so, mice that developed more severe compulsive behaviors were consistently convergent in microbiota responses (Figure 3 & 5) and the microbiota signatures reflected an alternative stable “disease” state dominated by pathobionts (Figure 6 and 7). Thus, even though correlation is not causation and the mechanism by which morphine-induced changes in the microbiota promotes or protects against compulsive behaviors have not yet been defined, this study paves the way for next steps. Interrogation of distinct genetic features, especially among the genera that significantly correlate with compulsivity score at both ends of the spectrum, could identify targets for study through genetic analysis. For those organisms that have yet to be cultured or are not amenable to manipulation, fecal microbial transplantation experiments could provide additional support and insight, especially if coupled with metagenomics, metabolomics and co-transcriptomic analysis of expression patterns of both microbiota and host. The role of local versus central nervous system for microbiota mediation or protection from compulsivity could be deciphered by targeted uncoupling such as with peripherally restricted antagonists. These are just some of the possibilities, any of which could provide new mechanistic insight into OUD development that reveals opportunities for disease intervention.

## CONCLUSION

Opioids remain essential for pain management but pose significant risks for the development of an OUD. Not all individuals exposed to opioids develop OUD, and an understanding of the underlying cause of this variability could provide important mechanistic insight into the process of progression of this chronic and relapsing condition, exposing new opportunities for prevention. Our findings demonstrate that distinct gut microbiota responses to prolonged morphine exposure correlate with compulsive drug-seeking. Beneficial bacteria linked to gut-microbiota-brain signaling that were relatively enriched in non-compulsive mice did so via dynamic association and flexible community interactions, likely related to functional redundancy, which could have facilitated their partial “recovery” of community structure post-morphine. In contrast compulsive mice developed a fragmented, pathobiont-driven microbiota that was unyielding post-morphine indicative of persisting dysbiosis. Even as it is not clear whether loss of beneficial taxa, or over-growth of pathogens, or both, contributed to progression of behaviors that develop due to neurobiological changes, which remain poorly defined and could vary from mouse to mouse, these studies do call for a more holistic understanding of gut-brain-microbiota interactions for understanding the role of the gut microbiota in OUD. They also highlight how leveraging natural protective mechanisms through microbiota manipulation could provide new avenues at mitigating the development of the debilitating disorder that is OUD.

## Supporting information

Supplemental Talbes

## ACKNOWLEDGMENTS

We thank Sarah W. Gooding, Madeline King, Joshua Gipoor, and Benjamin Wasson for assistance with data curation and collection; Maria Emanuel, Stephen Fiering, and Stephen Jones for their immeasurable support. We are grateful for the expertise of the staff at the Hubbard Center for Genome Studies at UNH and especially W. Kelley Thomas for advice, and Anthony Westbrook for bioinformatics assistance. We are grateful for the deep knowledge, helpful guidance, assistance and patience of Linnea Morley and Dean Elder at the Animal Resource Office at UNH.

## DATA AVAILABILITY STATEMENT

The 16S amplicons that support the findings of this study are available in NCBI under the accession number PRJNA1098090. The authors confirm that all processed data supporting the findings of this study are available within the article and/or its supplementary materials.

