## Supplemental Figures for "Dynamic flexibility of the murine gut microbiota to morphine disturbance enables escape from the stable dysbiosis associated with addiction-like behavior"

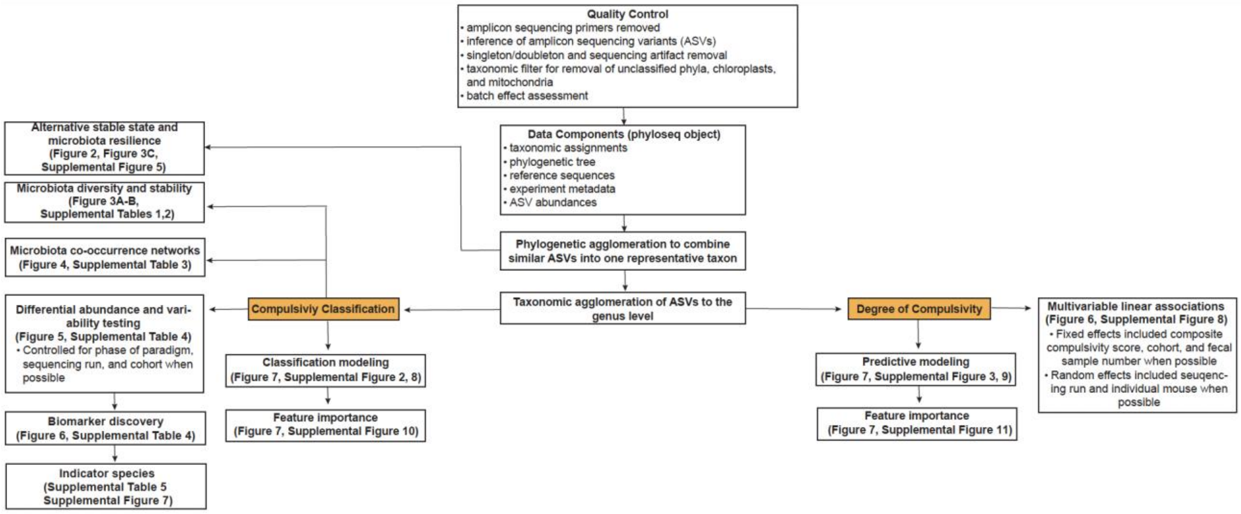


**Supplemental Figure 1: Pipeline of 16S rDNA sequencing read processing and subsequent analysis of amplicon sequence variants (ASVs) for associations with compulsivity classification and degree of compulsivity.** Demultiplexed raw sequencing reads were processed using cutadapt[41], DADA2[43, 44], and custom scripts for post-processing artifact removal to generate a quality-controlled dataset. Taxonomy was assigned using the GreenGenes reference database (v13.8)[45]. All data components (i.e., taxonomic assignments, experiment metadata, ASV counts, ASV sequences, and a phylogenetic tree) were stored together using the phyloseq package[48] for downstream analyses. Subsequent analyses utilized ASV datasets that were either agglomerated by phylogeny (similar ASVs into one representative taxon) or taxonomy (genus level) to investigate microbiota covariates with compulsivity classification ("compulsive" or "non-compulsive") and degree of compulsivity (composite compulsivity score). All analyses and visuals were produced using R Studio (v4.2.0).

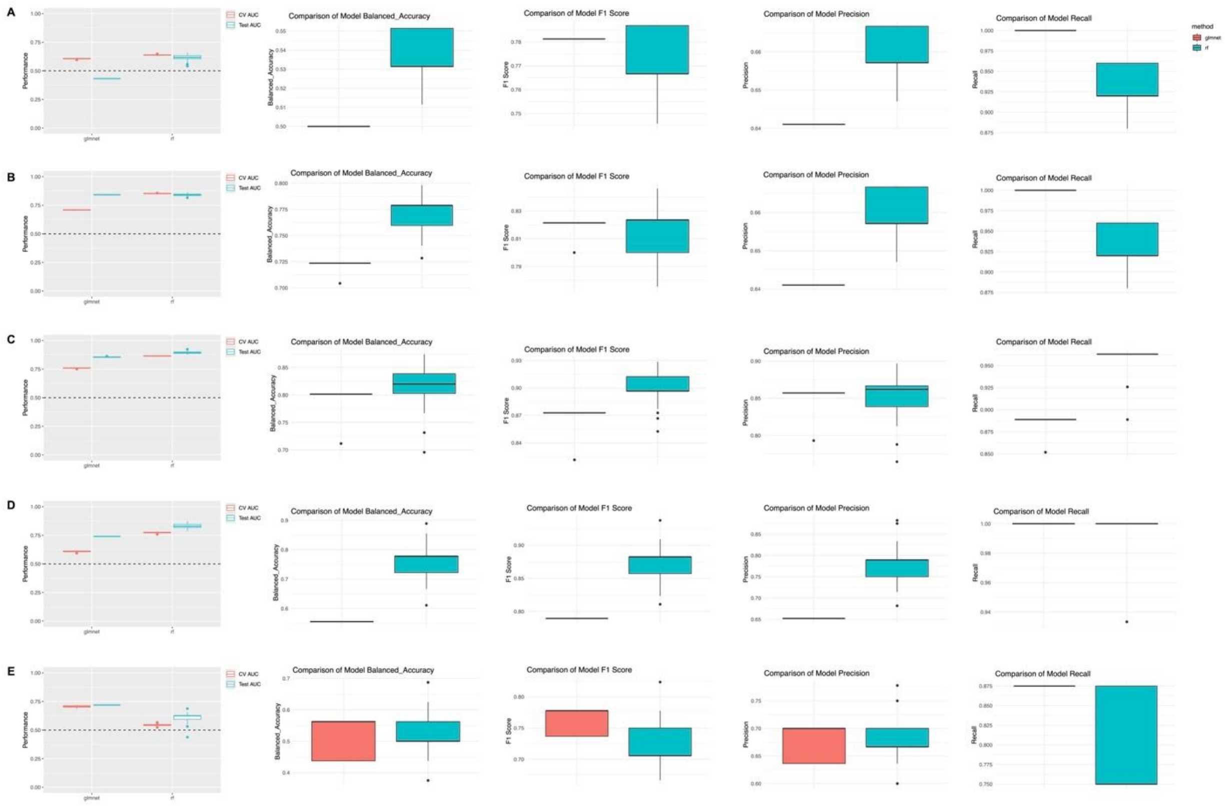


**Supplemental Figure 2: Comparison of performance metrics for model selection between L2 logistic regression and random forest algorithms for regression modeling.** L2 logistic regression and random forest models were trained on genus abundance counts to classify mice as compulsive or non-compulsive within individual phases of the paradigm using mikropml (v1.6.1)[74]. Case weights were calculated based on the proportion of compulsive and non-compulsive mice to address the imbalance between these groups (see methods for details). Phases of the paradigm included (A) pre-paradigm, (B) early morphine self-administration, (C) late morphine self-administration, (D) extinction, and (E) reinstatement and post-paradigm (combined for appropriate power to generate reliable models). Higher values in any performance metric indicate better model performance.

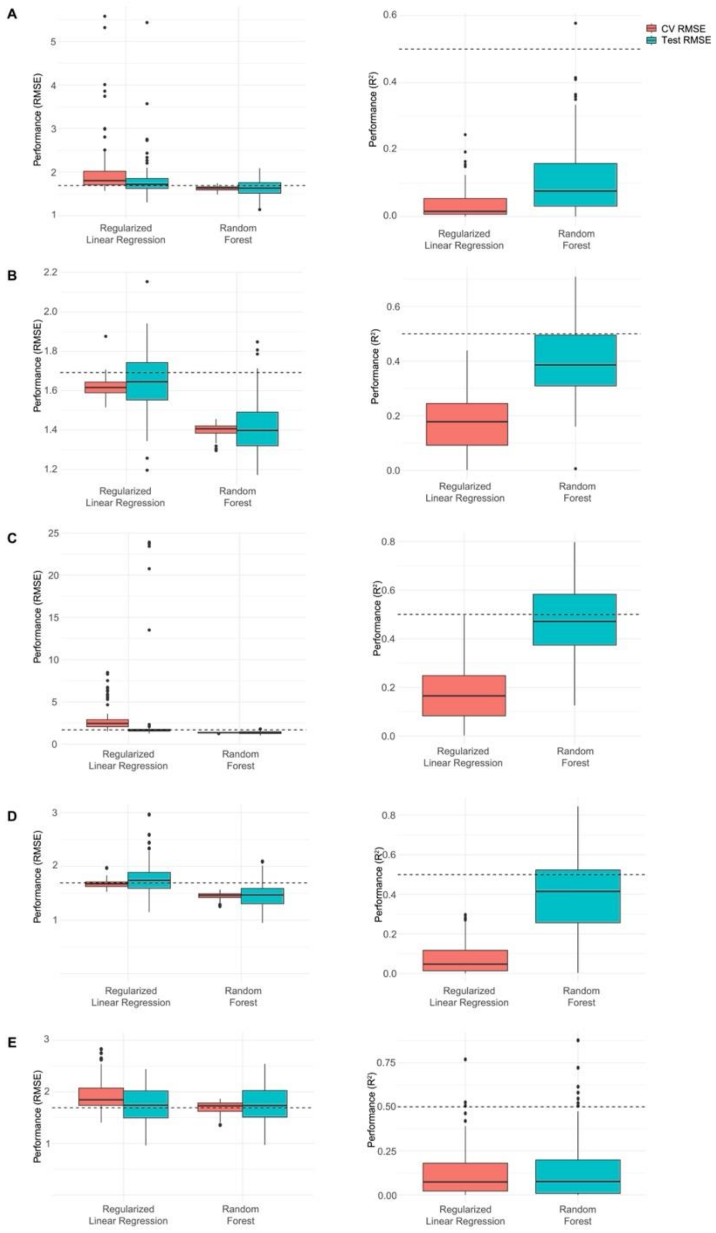


**Supplemental Figure 3: Comparison of performance metrics for model selection between L2 logistic regression and random forest algorithms for weighted classification modeling.** L2 logistic regression and random forest models were trained on genus abundance counts to predict community members most strongly associated with degree of compulsivity within individual phases of the paradigm using mikropml (v1.6.1)[74] (see methods for details). Phases of the paradigm included (A) pre-paradigm, (B) early morphine self-administration, (C) late morphine self-administration, (D) extinction, and (E) reinstatement and post-paradigm (combined for appropriate power to generate reliable models). Higher values in any performance metric indicate better model performance.

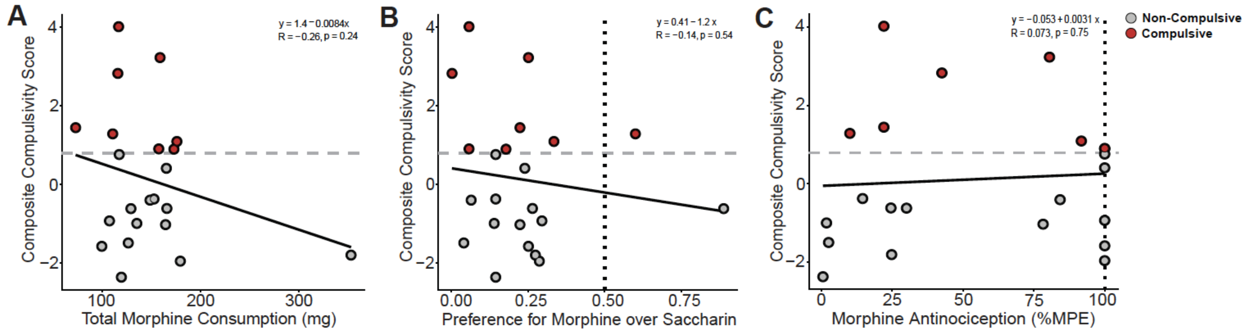


D

**Summary of behavior measures for individual mice**

| Classification | Mouse | Self-Administration Sub-Score | Extinction Sub-Score | Reinstatement Sub-Score | Composite Score | Total Morphine Consumption (mg) | Preference for Morphine Over Saccharin | Morphine Antinociception (%MPE) |
|---|---|---|---|---|---|---|---|---|
| Compulsive | 34M | 1.636322027 | 0.838277171 | 1.544490378 | 4.019089577 | 116.8 | 0.055555556 | 22 |
| Compulsive | 38M | 2.421802096 | 0.398493116 | 0.410414496 | 3.230709709 | 158.9 | 0.25 | 80.5 |
| Compulsive | 8M | -0.281641031 | 2.098006383 | 1.011104937 | 2.827470288 | 116 | 0 | 42.5 |
| Compulsive | 8F | -0.163902975 | 1.01190548 | 0.595765606 | 1.443768112 | 73.15 | 0.222222222 | 22 |
| Compulsive | 9M | 0.009467991 | 0.830199078 | 0.446324399 | 1.285991468 | 110.8 | 0.6 | 10 |
| Compulsive | 36M | -0.218051241 | 0.91447968 | 0.395177871 | 1.09160631 | 176 | 0.333333333 | 91.75 |
| Compulsive | 39M | 0.458346698 | -0.002683723 | 0.446938596 | 0.902601571 | 157.6 | 0.055555556 | 100 |
| Compulsive | 32M | 0.873067059 | -0.652398468 | 0.674649144 | 0.895317734 | 173 | 0.176470588 | 100 |
| Non-compulsive | 37M | 0.716561606 | -0.324843997 | 0.367204354 | 0.758921963 | 117.4 | 0.142857143 | 100 |
| Non-compulsive | 40M | -0.073561494 | 0.572711306 | -0.090976404 | 0.408173408 | 165.3 | 0.238095238 | 100 |
| Non-compulsive | 41M | 0.855862571 | -0.85536279 | -0.376142859 | -0.375643078 | 152.7 | 0.142857143 | 14.5 |
| Non-compulsive | 29M | 0.13830296 | -0.666714978 | 0.124089324 | -0.404322695 | 148.7 | 0.0625 | 84.25 |
| Non-compulsive | 33M | 0.093903303 | -0.422985587 | -0.286970939 | -0.616053224 | 165.8 | 0.263157895 | 24.5 |
| Non-compulsive | 6F | -0.36399855 | -0.553093221 | 0.296426177 | -0.620665594 | 129.3 | 0.888888889 | 30 |
| Non-compulsive | 30M | -0.13186301 | -0.38054046 | -0.420382182 | -0.932785652 | 107.5 | 0.294117647 | 100 |
| Non-compulsive | 6M | -0.474346476 | -0.307971614 | -0.215280868 | -0.997598958 | 135.3 | 0.137931034 | 1.75 |
| Non-compulsive | 27M | 0.18643341 | -0.424053881 | -0.791408026 | -1.029028497 | 164.5 | 0.222222222 | 78.25 |
| Non-compulsive | 26M | -0.664531076 | -0.329000115 | -0.502371307 | -1.495902498 | 126.6 | 0.038461538 | 2.5 |
| Non-compulsive | 31M | -0.591023956 | -0.490535108 | -0.499870811 | -1.581429876 | 99.7 | 0.25 | 100 |
| Non-compulsive | 28M | -0.81008659 | -0.446732717 | -0.548513849 | -1.805333156 | 352.9 | 0.272727273 | 24.75 |
| Non-compulsive | 7F | -0.89363969 | -0.723245801 | -0.342516946 | -1.959402437 | 179.5 | 0.285714286 | 100 |
| Non-compulsive | 35M | -0.811932905 | -0.959196576 | -0.597726244 | -2.368855725 | 119.6 | 0.142857143 | 0.5 |

**Supplemental Figure 4: Composite compulsivity score does not correlate with degree of morphine consumption, morphine preference, or antinociceptive tolerance to morphine.** Linear regression analyses of (A) total morphine consumption (y = 1.4 - 0.0084x; R2 = -0.26; Pearson's correlation p-value = 0.24), (B) preference for morphine over saccharin (y = 0.41 – 1.2x; R2 = -0.14; Pearson's correlation p-value = 0.54), and (C) degree of antinociceptive tolerance (y = -0.053 + 0.0031x; R2 = 0.073; Pearson's correlation p-value = 0.75) with composite compulsivity scores. Horizontal dashed light grey lines represent the population average + 1 interquartile deviation (IQD) used for compulsivity classification, where mice with scores above the dashed line were classified as "compulsive" drug-seekers, and mice with scores below the dashed line were classified as "non-compulsive" drug-seekers. The vertical dotted black line represents the level of morphine preference over saccharin (<0.50 = lower preference; >0.50 = higher preference) in (B), or the development of antinociceptive tolerance (100% MPE = non-tolerant; <100% MPE = tolerant) in (C). Summary of behavior measures for individual mice are detailed in (D) in descending order of composite compulsivity score. The bolded grey dashed line indicates the threshold for compulsivity classification (population average + 1 IQD). The bolded black dashed lines indicate the threshold for mice with extreme compulsivity classification (population average +/- 2 IQD).

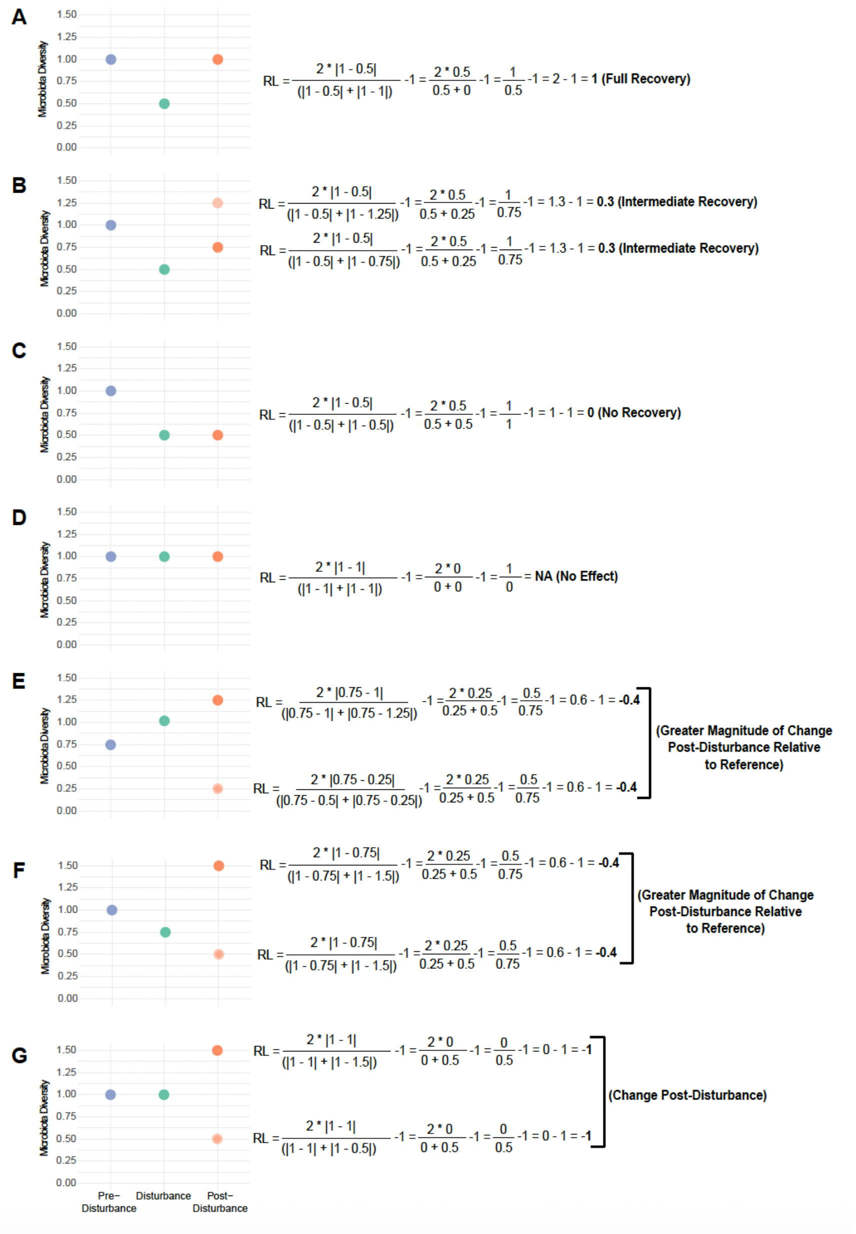


**Supplemental Figure 5: Examples of how aspects of microbiota diversity may impact the interpretation of the trajectory of recovery from scores generated using the RL index formula.** A) An RL of "1" indicates the complete recovery post-disturbance (a return to the reference state, ending diversity = starting diversity). B) Positive values between "0" and "1" represent intermediary recovery. C-D) An RL of "0" indicates no recovery (no change from the disturbed state to the post-disturbance state, ending diversity = disturbed diversity). E-F) Negative values between "0" and "-1" indicate a greater relative change in diversity of the recovery microbiota than disturbance microbiota. G) An RL of "-1" indicates that no change occurred during disturbance (starting diversity = disturbed diversity).

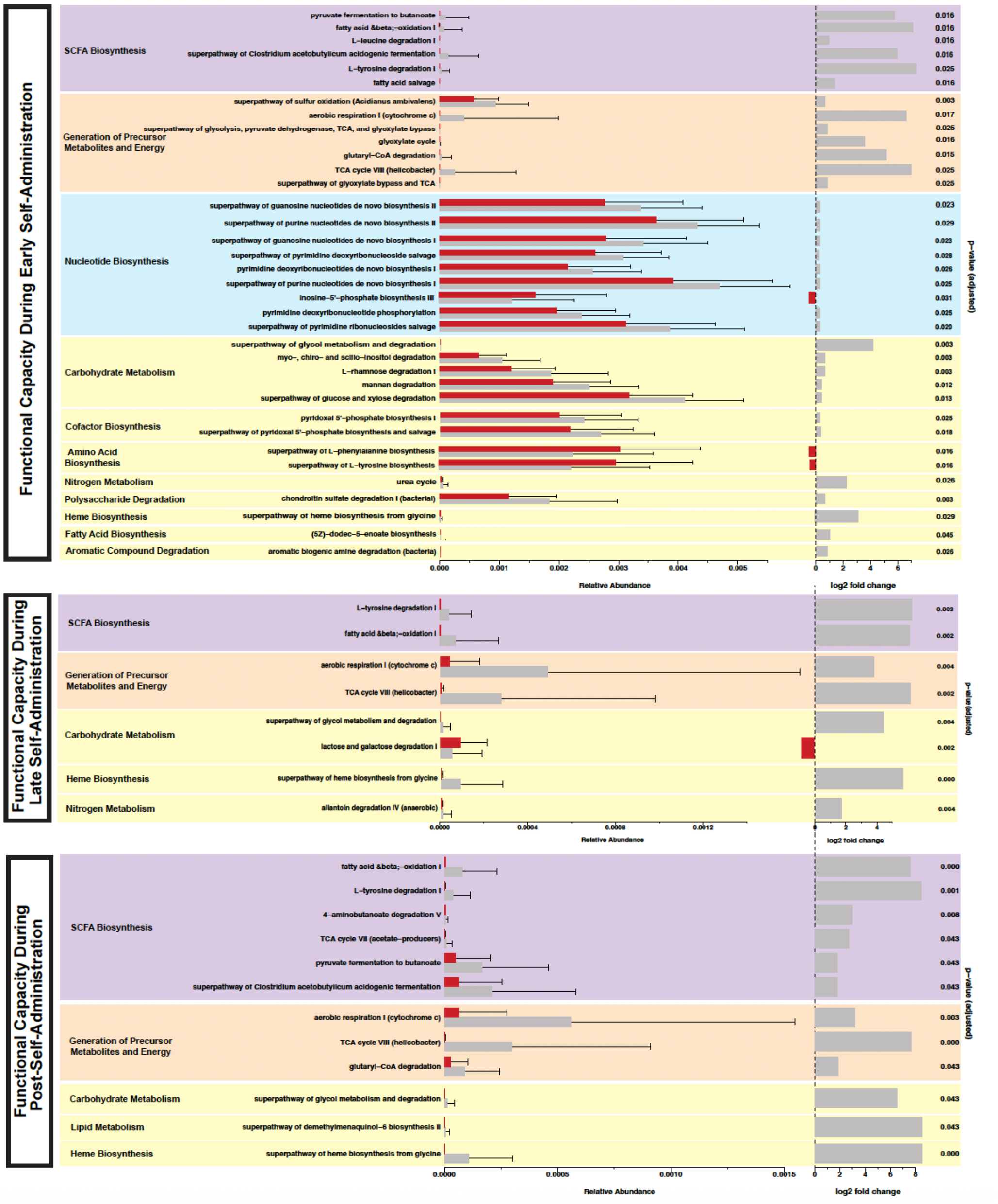


**Supplemental Figure 6: Differences in predicted functional capacity between compulsive and non-compulsive mice during and post-morphine exposure.** Predicted functional profiles (metabolic pathways and abundances) derived from 16S rDNA sequences were generated using the PICRUSt 2.0 pipeline[60]. Differential abundance testing of predicted metabolic pathways between compulsive (red) and non-compulsive (grey) mice was compared using the LinDA model using the ggpicrust2 (v.1.7.3) package[66] in RStudio (v.4.2.0).

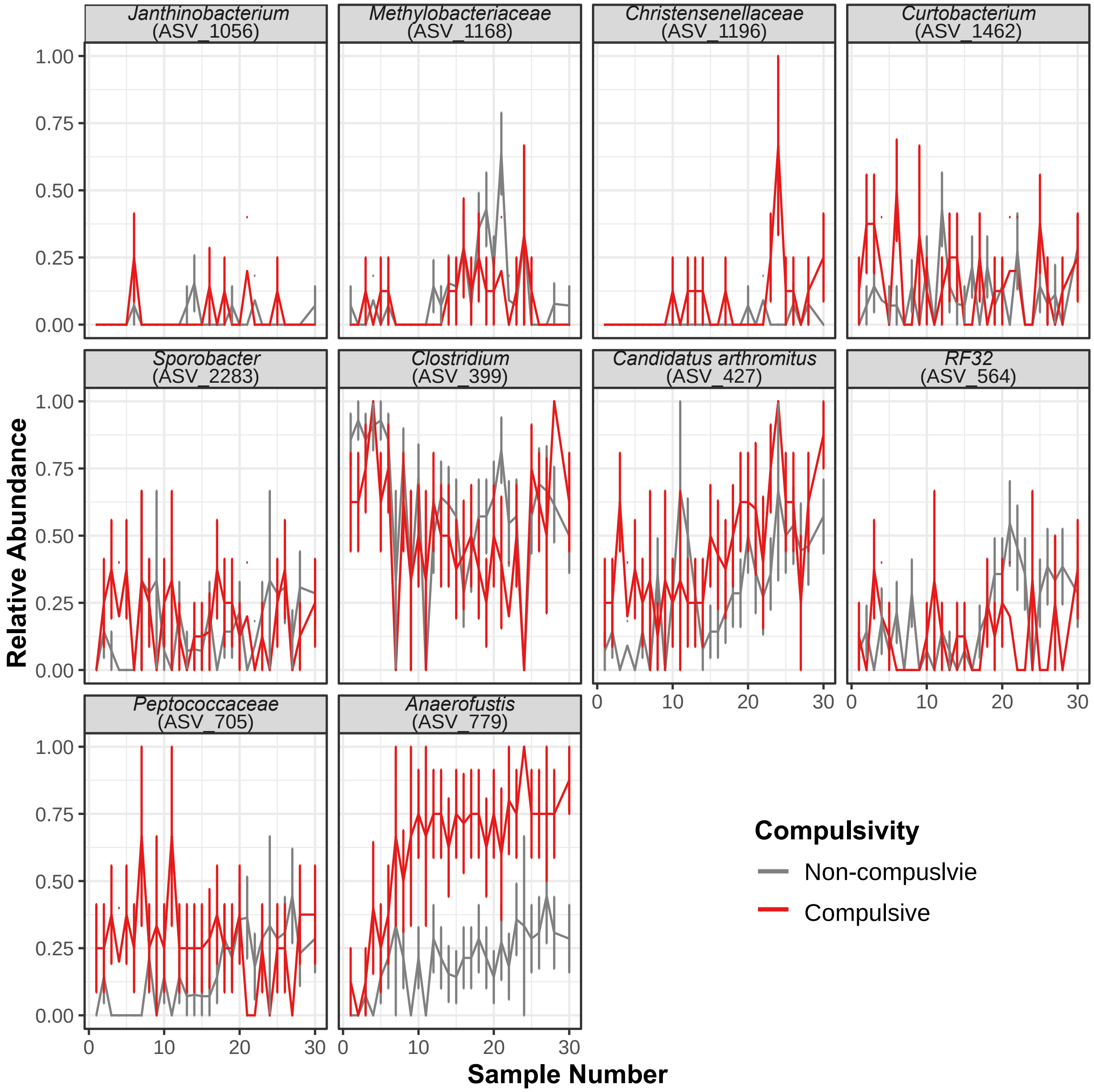


**Supplemental Figure 7: Temporal patterns of indicator species identified in non-compulsive and compulsive mice.** Average relative abundances of genera identified as indicators using IndicSpecies package in R[71] of non-compulsive (grey) and compulsive (red) mice where sample numbers represent the consecutive number of samples as a proxy for time and error bars represent standard error.

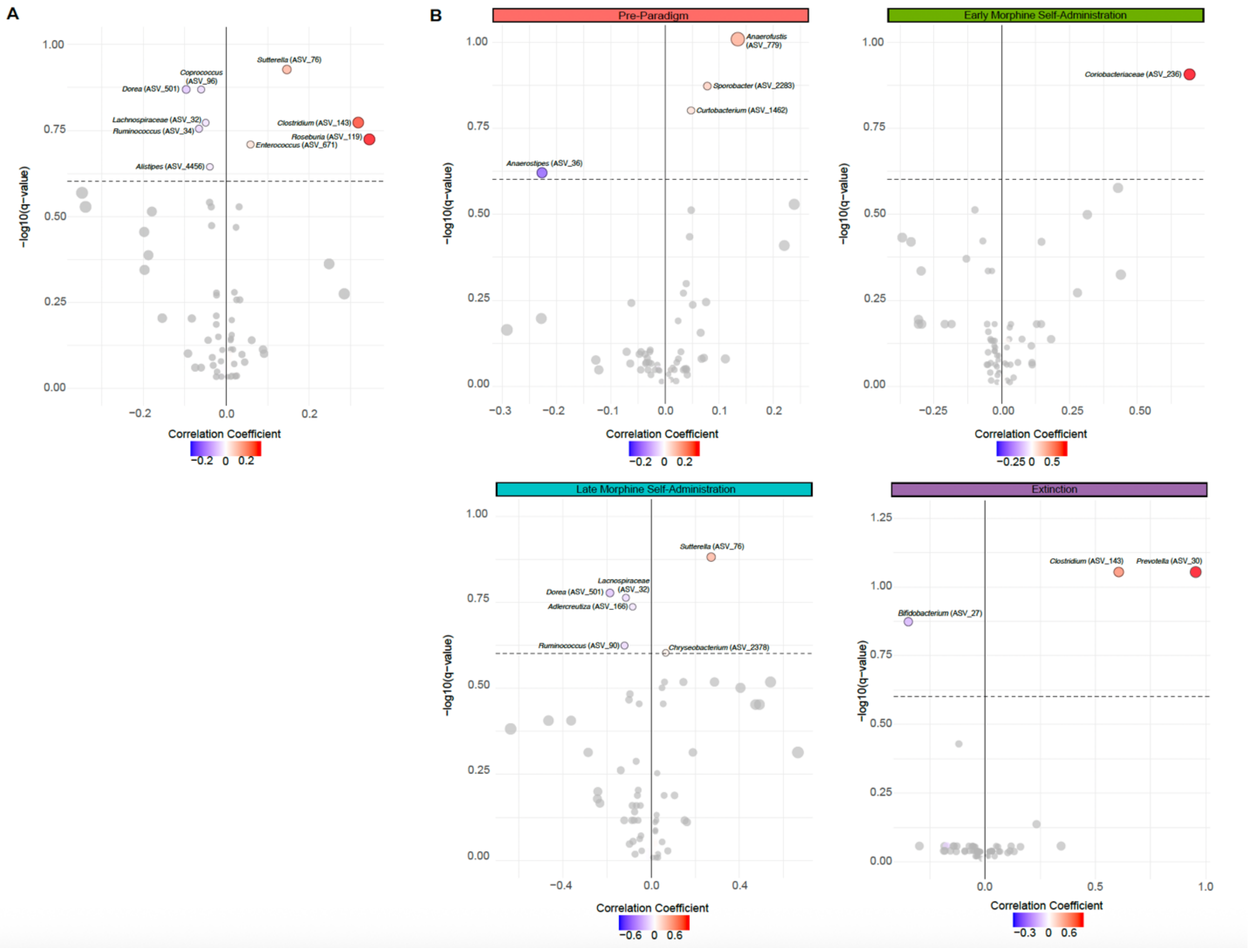


**Supplemental Figure 8: Gut microbiota genera correlated with composite compulsivity scores across all phases of the paradigm.** Associations of center-log ratio (CLR) transformed relative abundances of individual genera (labeled as ASV) with composite compulsivity score from all phases combined (A) or using microbiota from a given stage or phase (B) were assessed using multivariable association testing with linear models (MaAsLin 2)[72]. Genera identified as significantly correlated based on the default threshold (q-value of 0.25, noted as the inverse log q = 0.6) are labeled and colored circles above the black dashed line and genera not significantly correlated are unlabeled grey circles below the black dashed line. The size of individual circles corresponds to the magnitude of effect size, or the strength of the correlation. Negative correlation coefficient values indicate inverse correlations with composite compulsivity scores, and positive correlation coefficient values represent direct correlations with composite compulsivity scores distributed horizontally. No genera significantly correlated with composite compulsivity scores during the reinstatement phase nor post-paradigm.

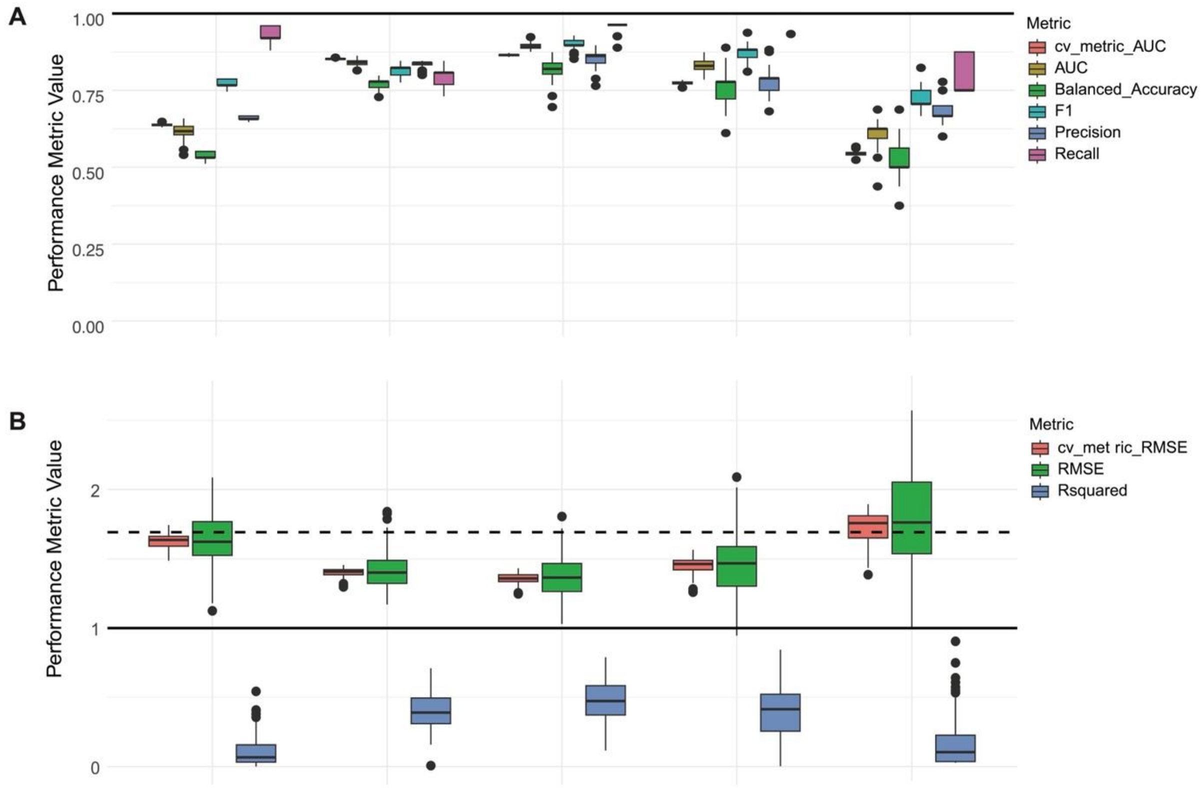


**Supplemental Figure 9: Evaluation metrics for performance of random forest models using categorical assignments or continuous composite scores during each phase of the paradigm.** A) Relevant metrics for model performance (suitable for an imbalanced dataset with more non-compulsive than compulsive mice) (bottom panel) are also shown. Higher metric values indicate better model performance at each phase (see methods for details). An AUC closer to 1 indicates stronger discrimination between groups. Balanced accuracy, which averages sensitivity and specificity, is well-suited for imbalanced sample groups, with values closer to 1 indicating high performance across both classes. The F1 score balances precision and recall, making it ideal for assessing performance on the minority class without bias from the majority class, with values closer to 1 reflecting strong performance in identifying true positives and minimizing false positives and false negatives. Precision and recall were also evaluated separately, with precision measuring the proportion of true positives out of all predicted positives and recall measuring the proportion of true positives out of all actual positives. Values closer to 1 for both precision and recall indicate high performance in predicting true positives accurately B) Relevant metrics for model performance (suitable for continuous data) (bottom panel) are also shown. Metric values below the baseline RMSE model (black dashed line) indicate better model performance relative to predicting the mean composite compulsivity score (see methods for details). Lower RMSE values indicate better predictive accuracy. Within the context of our data, a baseline RMSE model was calculated from the composite compulsivity score mean as a reference point for further evaluation of a model's performance. An RMSE value lower than the baseline RMSE value indicates improvement beyond the mean-prediction model. R-squared values show the variance explained by genus abundances, with values closer to 1 indicating higher explanatory power.

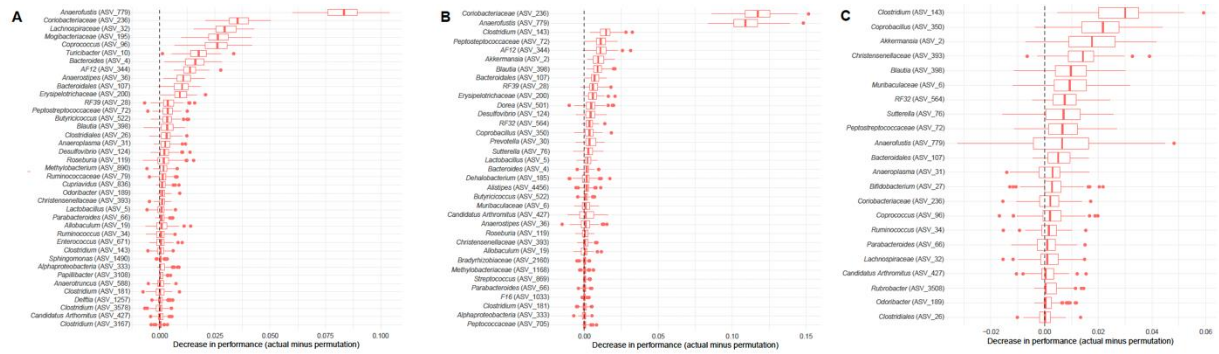


**Supplemental Figure 11: Total number of features identified as important contributors to regression model performance using random forest algorithms.** Feature importance of genera at each phase of the paradigm was assessed using permutation testing in mikropml (v1.6.1)[74] (see methods for details). Phases of the paradigm included (A) early morphine self-administration, (B) late morphine self-administration, and (C) extinction). Features which have a lower performance when permuted have a difference in performance above zero. The features with the greatest decrease are the most important for model performance.

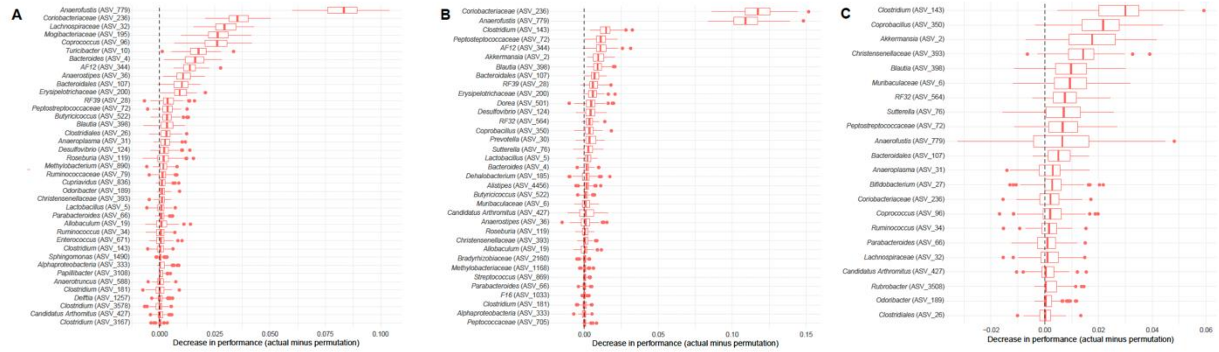

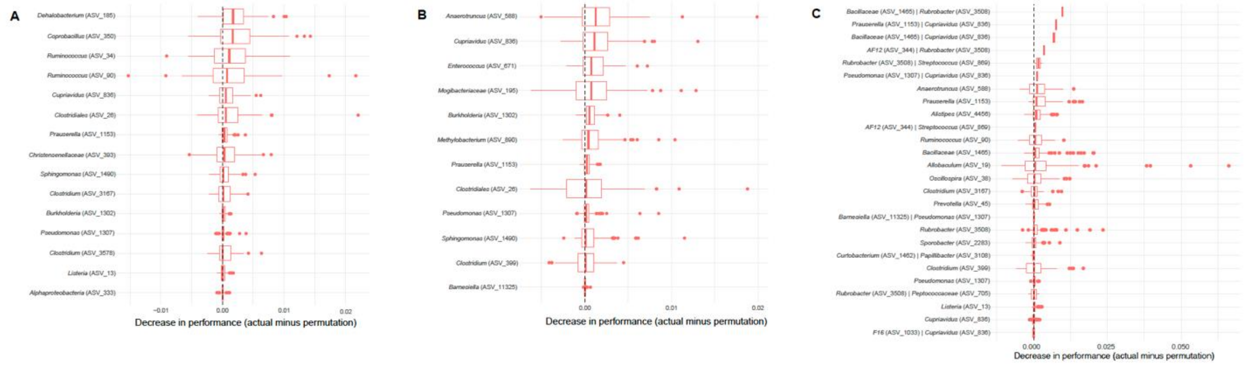


**Supplemental Figure 11: Total number of features identified as important contributors to regression model performance using random forest algorithms.** Feature importance of genera at each phase of the paradigm was assessed using permutation testing in mikropml (v1.6.1)[77] (see methods for details). Phases of the paradigm included (A) early morphine self-administration, (B) late morphine self-administration, and (C) extinction). Features which have a lower performance when permuted have a difference in performance above zero. The features with the greatest decrease are the most important for model performance.
